# Elucidating the perception, awareness and biosecurity practices among the live poultry/poultry-meat sellers to mitigate the risk of zoonotic salmonellosis

**DOI:** 10.1101/2025.06.17.660267

**Authors:** Mirza Mienur Meher, Marya Afrin, Abdullah Al Bayazid, Nashib Parajuli, Md Sohel Rahman, Md Sayedul Islam, Md Zulfekar Ali, Md. Mominul Islam

## Abstract

Zoonotic salmonellosis remains a major public health concern, especially in low-resource settings where poultry handling practices are often informal. Biosecurity measure of live poultry/poultry-meat vending shop is necessary to address this zoonoses. This study investigated the knowledge, attitudes, and biosecurity practices of poultry/poultry-meat sellers toward the risk of zoonotic salmonellosis in Bangladesh. A quantitative, descriptive, cross-sectional study was conducted among the 432 poultry/poultry-meat sellers through a structured questionnaire in multistage sampling approach. Majority of the poultry/poultry-meat sellers (n=432) were male (75.5%), age between 26-34 years (31.9%), had primary education (28.9%), experience of ≤5 years (40.3%), and monthly income was <20000 BDT (43.3%). Additionally, minimal had training on live poultry marketing (33.1%), and poultry-meat processing and marketing (28.5%). The 33.34% and 62.03% of poultry/poultry-meat sellers had the good knowledge and positive attitudes respectively. While 41.43% were involved in correct biosecurity practice. Moreover, the mean KAP score was 53.12±0.89, 68.59±0.79 and 54.58±0.73 respectively. From the multivariate analyses, higher education was a strong predictor of good knowledge (Bachelor’s degree: OR=6.64, 95%CI:2.59-17.04, p<0.01), positive attitude (Bachelor’s degree: OR=13.48, 95%CI:3.72-48.87, p<0.01), and correct biosecurity practices (HS: OR=5.52, 95%CI:2.62-11.63, p<0.01; Bachelor’s degree: OR=4.81, 95%CI:2.03-11.39, p<0.01). The sellers with good knowledge, positive attitudes, and correct biosecurity practices perceived higher zoonotic risk (median=7, p<0.05), with perception co-relation that significantly influenced by education (r=0.131, p<0.01) and training (r=0.084, p<0.05). Despite moderate awareness, sellers showed notable gaps in safe practices, highlighting the need for practical training that enhances risk perception, promotes behaviour change, and builds hands-on biosecurity skills.

## Introduction

Zoonotic diseases remain a persistent public health threat globally, especially in the regions where intensive human and animal interactions, informal food markets, and inadequate hygiene practices are common. Zoonotic illnesses have a twofold impact on people’s well-being since they endanger the health and output from the poultry or animals. Almost 60% of recognized infectious illnesses are zoonotic, as are up to 75% of emerging infectious pathogens [1]. A large number of zoonoses are regarded as risks to occupational health [2]. Live poultry handlers are at always risk because different kinds and intensities of interaction may lead to outbreak of zoonoses. The poultry or meat handler working in slaughterhouses can also be an asymptomatic reservoir of food borne microorganisms as well as zoonoses [3]. Among the most prevalent foodborne zoonotic pathogen, Salmonella isolates are the major that can be transmitted through the live poultry or poultry meat to humans [4,5]. Globally, non-typhoidal *Salmonella* infections cause an estimated 93.8 million food-borne illnesses and 155,000 deaths annually, due to the food contamination [4]. The non-typhoidal Salmonella*, Salmonella enteritidis* and *Salmonella typhimurium* are the most prevalent serovars of Salmonella that associated with poultry and pose a risk to human (Thorns, 2000; Wang et al., 2020). Worldwide, salmonella infections pose a serious threat to public health and cause financial hardship because of the high costs of illness prevention and treatment. [6]. Particularly, poultry harbour Salmonella in their gastrointestinal tract and excrete the pathogen through faeces, contaminating the environment [5]. During processing of poultry meat, cross-contamination can occur at multiple points, including slaughter, evisceration, and packaging [5]. Though the consumption of contaminated poultry meat and eggs is the primary source of human salmonellosis [7], the live poultry/poultry-meat handler also possess risk due to direct contamination. The environmental alterations caused by climate change significantly influence the survival and transmission of food born zoonotic pathogen, Salmonella [8]. This underscores the importance of the One Health approach, which recognizes the interconnectedness of human, animal, and environmental health in understanding and mitigating such emerging public health threats.

One of the major and fastest-growing subsectors in Bangladesh, poultry has created a significant amount of job opportunities and is essential to reducing poverty and ensuring food security [9–13]. Live poultry markets (LPM) for vending the poultry meat by dressing, processing in open environments are common practice in Asia, including Bangladesh [14,15]. These LPM ranges from the wet markets to informal roadside markets [15]. Basically, the sellers at LPM offer live poultry for consumer inspection and selection, usually offering slaughtering services on-the-spot that enable consumers to witness the poultry slaughtered to confirm they get meat from live poultry [15]. These LPM are recognized hotspots for the spread of zoonotic pathogen from live poultry to humans creates public health risks [15,16]. Live poultry vendors are usually involved in high-contact activities with little understanding or training in disease control, and these environments often lack official regulatory supervision. Biosecurity in these environments is frequently compromised by overcrowding, poor waste disposal, inadequate sanitation, and lack of protective equipment which increase the risk of pathogen persistence and transmission [17]. While many studies across the world have focused on knowledge, attitude and practices on food safety [14,18–23], and other zoonoses [24,25,25,26], the KAP study of live poultry/poultry-meat sellers on zoonotic salmonellosis almost remain unexplored. Understanding the KAP of these frontline workers is essential for identifying risk perceptions and behavioural gaps which may contribute to ongoing transmission of zoonoses. Existing research highlights that knowledge and positive attitudes significantly influence hygienic practices, but these relationships vary with educational level, socioeconomic conditions, and access to training [14,21,23,27,28]. However, evidence from informal poultry markets in South Asia remains limited. There is a pressing need to evaluate whether poultry sellers in these environments comprehend the zoonotic risks posed by *Salmonella*, and whether their practices reflect this awareness. Although certain training initiatives and interventions have been investigated, little is understood about how demographic characteristics like education, income, gender, or training background impact sellers’ willingness to take part in disease prevention programs, their perceptions of risk, and their confidence in putting biosecurity measures in place. Filling these gaps is critical for developing culturally and contextually relevant education and intervention programs. Therefore, evaluation of biosecurity measure of live poultry/poultry-meat vending shop is necessary to address the increasing incidence of foodborne zoonoses and design potent strategies to refine meat hygiene and public health. As far as our observation goes, this is for the first time ever study of poultry/poultry-meat seller’s knowledge, attitude and practices on zoonotic salmonellosis in Bangladesh. In this present study we aimed to investigate the knowledge, attitudes, and biosecurity practices of poultry/poultry-meat sellers toward the risk of zoonotic salmonellosis in Bangladesh.

## Materials and methods

### Ethical Approval and Participant Consent

The ethics of this study was in accordance to research ethics followed by the Department of Microbiology and Public Health of this respective University. The ethical consent number was BSMRAU/FVMAS/MPH/20(Ethical Approval)/2020/06; Date: 07-11-2021. Prior to data collection, all participants were informed about the purpose, procedures, confidentiality, and voluntary nature of their participation. Verbal informed consent was obtained from each participant. The data collector documented this consent on the questionnaire form and recorded the name of a witness present during the consent process. No second member of the research team was required to serve as a witness. In the case of minors, consent was obtained from a parent or legal guardian. The study involved no clinical procedures, no collection of biological samples, and posed minimal risk to participants. All data were anonymized prior to analysis to ensure participant confidentiality, and no information that could directly identify individual participants was available to the authors during or after data access.

### Study area

This study was conducted in 12 selected districts of (Mymensingh, Kishoreganj, Netrakona, Jamalpur, Tangail, Sherpur, Dhaka, Narayanganj, Gazipur, Manikgonj, Munshigonj, Narsingdi) Bangladesh, particularly, the central part of Bangladesh [29]. The study area is located between 24°340′ to 23°810′ north latitude and 89°400′ to 90°980′ east longitude. The region has an average annual rainfall of about 1600mm, with the average environmental temperatures of about 25°C [29]. This study area, encompassing the high concentration of poultry farms and live poultry markets [9,30], which play a crucial role in the country’s poultry supply chain. Additionally, the region’s dense population and significant poultry trade activities increase the risk of zoonotic disease transmission, making it a vital area for understanding biosecurity practices. The spatial location of the study areas where data was collected shown in Fig 1. The ArcGIS-ArcMap version 10.8 (ESRI, USA) software was used to prepare the study map.

**Fig 1.**
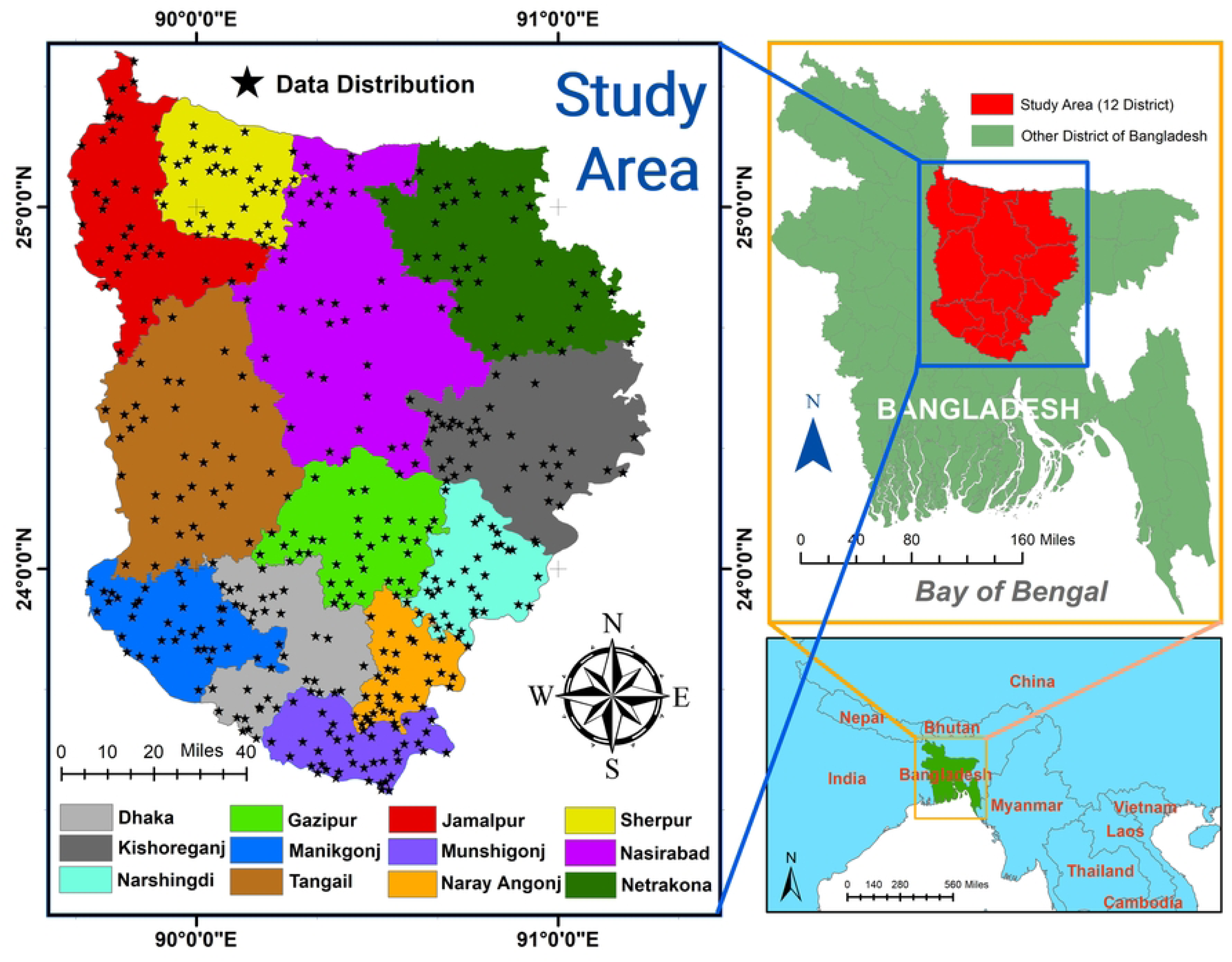
Spatial location of the study population (live poultry/poultry-meat sellers) in different area of Bangladesh.

### Study Design and Sampling methods

A quantitative descriptive cross-sectional study was conducted among the live poultry/poultry-meat sellers in the selected area of Bangladesh from 01 July 2021 to 30 September 2023 by using a structured pretested questionnaire to quantify the knowledge, attitude, and biosecurity practices of poultry/poultry-meat sellers. were interviewed face to face during marketing hours without prior notice of the interview. The purpose of the study was briefed to the poultry/poultry-meat sellers before the questionnaire and the poultry/poultry-meat sellers were assured about the confidentiality of their status. The questionnaires were translated from English to the mother tongue “Bengali” and the responses were recorded into the English language. And finally completed by an interviewer in individual interviews. The respondents were given sufficient time to answer the questionnaire. In this study, percipients were eligible to participate according to specific inclusion/exclusion criteria (S1 Appendix). A multistage sampling technique was used to ensure representativeness across the central region of Bangladesh. First, a simple random sampling method was applied to select the poultry markets within each district. Then, the participants (poultry/poultry-meat sellers) were selected using systematic random sampling based on their availability and willingness to participate.

### Sample Size Determination

The required sample size for this study was estimated {eq. (1)} according to the formula [31] with a little modification

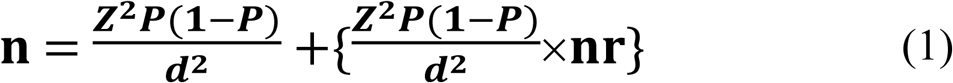

Where,

*n* = sample size,

*Z* = 1.96 (95% confidence level),

*P* = expected prevalence or proportion (in proportion of one; when good knowledge is 53.33%,

*P* = 0.53; when positive attitude is 80.00%, *P* = 0.80 and when correct practice is 30.00%, *P* = 0.30),

*d* = precision (in proportion of one; whereas *P*=0.1 to 0.9, therefore *d* = 0.05), and nr = non-response rate (if it was assumed that 15% of calculated sample may not response, then 15% of total sample will be increase size; nr = 0.15)

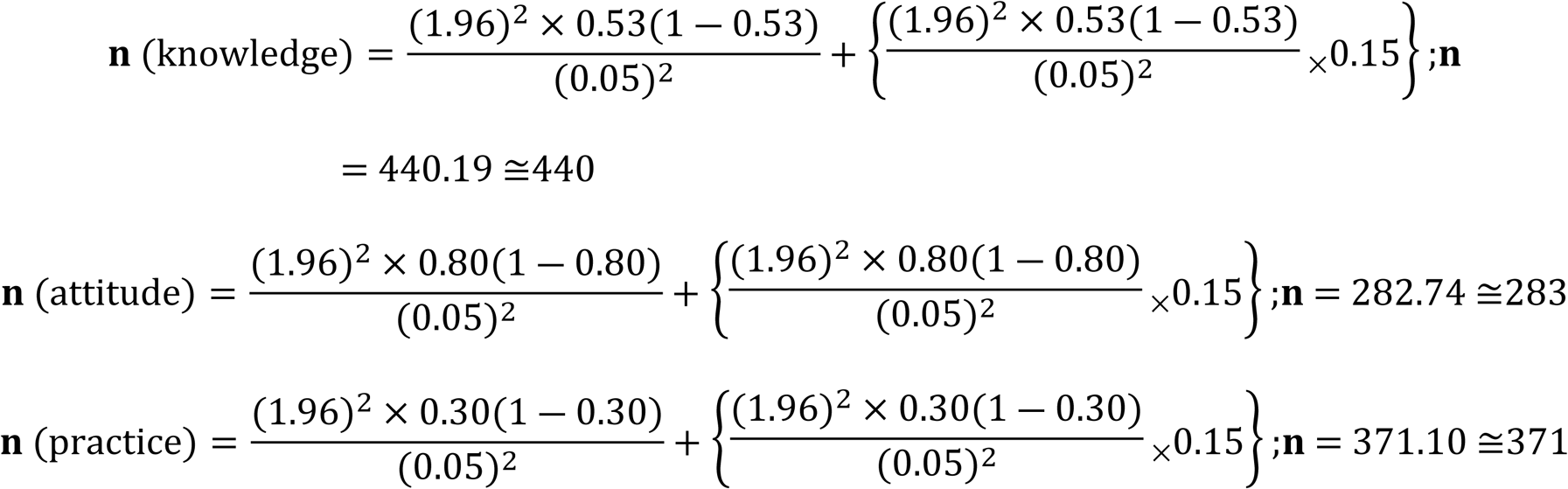

The expected proportion of poultry/poultry-meat sellers having good knowledge, positive attitude, and correct practice were 53.33%, 80.00%, and 30.00 respectively, obtained through a pilot study of a small group of poultry/poultry-meat sellers (n=30). The calculated sample size for knowledge, attitude, and practice were 440, 283, and 371. Among the three calculated sample size, the highest value (440) was considered as expected respondent. And finally, a total of 432 poultry/poultry-meat sellers willingly responded, were considered as sample size in this study.

### Questionnaire

A structured questionnaire was used to assess the zoonotic salmonellosis knowledge, attitudes and practices of poultry/poultry-meat sellers, were adapted from similar type past studies [14,21,23,27,32]. The questionnaire consists of four sections. Section 1 comprised questions relating to socio-demographic characteristics, including sex, age, educational qualifications, experience, monthly income, and training on live poultry marketing and poultry meat processing. Section 2 comprised 15 close-ended questions on poultry/poultry-meat sellers knowledge questions relating to poultry/poultry-meat sellers’ knowledge on zoonotic salmonellosis, including the zoonoses, salmonellosis as a zoonotic disease, food poisoning caused by salmonella, zoonotic salmonella transmission, contamination of dressed poultry meat by zoonotic Salmonella spp., risk for zoonotic salmonellosis, washing and cleaning, use of disinfectant, carcass dressing water, and disposal of poultry wastages. Each question was provided by three possible answers (yes, no and I don’t know). Section 3 included the questions about the zoonotic salmonellosis attitude of poultry/poultry-meat sellers which comprised 13 questions including the perspectives on the importance of worker training, proper hygiene and sanitation, avoiding slaughtering sick birds, maintaining a clean environment during meat dressing, washing with safe water, proper disposal of poultry offal, preventing contamination with feathers or droppings, cooking meat thoroughly, avoiding handling raw meat with wounded hands, refraining from touching the face or hair during processing, using gloves and masks, maintaining short and uncolored nails, and avoiding handling money while processing raw meat. Each questions had the seven possible answers (Strongly agree, moderately agree, partially agree, partially disagree, moderately disagree, strongly disagree, and neutral). Lastly, the section 4 included 19 questions assessing self-reported biosecurity practices such as separating poultry by age and type, restricting food, drink, and smoking in the shop, using fresh and hot water for cleaning and dressing, washing hands with soap before and after meat processing, cleaning and washing utensils, and disinfecting the shop. Additionally, questions covered avoiding jewelry, wearing protective gear (masks, aprons, caps, gloves), changing clothes post-work, and proper disposal of slaughtering waste, including blood, viscera, feathers, and dead birds. Measures to control pests and restrict wild birds, dogs, and cats near the shop were also assessed. Each questions had the four possible answers (always, often, rarely, and never).

### Measures

Each question of the section 2 (question for assessment of knowledge) of structured questionnaire had three possible answers, where the allocated point was 1 for “yes” and 0 for “no” and “don’t know” [dog cat paper]. The possible score range for knowledge section was 0–15 points. Then the scores were converted to 100 points. In the third section of the questionnaire, the allocated points for the answer (strongly agree=6; moderately agree=5; partially agree=4; partially disagree=3; moderately disagree=2; strongly disagree=1 and neutral=0) of each question were used to evaluate the poultry/poultry-meat sellers’ attitudes. The score range was between 0 to 78 and converted to 100 points. In the section four, the questions regarding the biosecurity practices, where the point 3, 2, 1, and 0 was awarded for the possible response of “always”, “rarely”, “often”, and “never” respectively. The possible score range for practices section was 0–57 points. After that the scores were converted to 100 points. Subsequently, ≥ 66.66 points out of 100 (two thirds) were considered as cut-off points for good knowledge, positive attitude, and correct biosecurity practices. Following this, those falling below the specified threshold was categorized as having poor knowledge, negative attitudes, and incorrect practices. Lastly, a Likert scale of 0 to 10 were applied for the self-assessment of perceived, confidence and interest of poultry/poultry-meat sellers toward the zoonotic salmonellosis.

### Validity and reliability of questionnaire

Prior to conduct the final study, a pilot study was performed among a randomly-selected small group of poultry/poultry-meat sellers (n=30) to determine the fitness of the questionnaire. The internal consistency reliability for the poultry/poultry-meat seller’s knowledge, attitudes and practices is judged based on the average interitem correlation (AIC) analysis and calculating Cronbach’s alpha. The AIC was 0.16 (−0.40 to 0.97), 0.38 (−0.17 to 0.70) and 0.12 (−0.49 to 0.89) for knowledge, attitudes and practices sections. The internal consistency (Cronbach’s alpha) was 0.710, 0.886 and 0.729 for knowledge, attitudes and practices sections of the questionnaire, indicating acceptable reliability [21,33]. The results of the piloted survey were not comprised in this final study.

### Statistical analysis

The data achieved from the survey were inputted and arranged accordingly into a spread sheet of Microsoft Excel-2019 and subsequently transferred into Statistical Package for Social Sciences (SPSS) version 27.0 for further analysis. Descriptive statistics was applied to calculate the frequencies, percentages, mean, standard error of mean, and KAP scores for each socio-demographic variable of the respondent’s and presented in tables. Pearson’s chi-squared test was used to test the association between categorical variable. Particularly, when more than 20% of cells of 2×2 contingency table had expected count less than 5, the p value of continuity correction was considered but when the table more than the 2×2 contingency then p value of Fisher exact tests was accounted. Additionally, Independent samples t-testing was conducted to compare the mean of KAP score of two community variables such as sex, TLPM and TPMPM. On the other hand, the one-way ANOVA was considered, followed by Tukey’s test, to compare the mean of KAP scores for demographic variable having more than two category level such as age, education, experience, and monthly income. The Binary logistic regression analysis with a 95% confidence interval was performed to evaluate the associations to determine associations between independent (sociodemographic variable) and dependent (KAP) factors. At first the Univariable logistic regression analysis was performed and then only univariable factors with p ≤ 0.05 were included for the multivariable logistic regression analysis. The Pearson correlation test (r) was performed to ascertain the strength of the correlations within the KAP scores. The strength of the correlations was classified as followed by another research [34]. The Mann– Whitney test was used to assess the differences within the distribution of ordinal variables expressed by 1 to 10 Likert scale in boxplot among the KAP level of the participants. Lastly, the Kendall’s tau-b rank correlation was considered to assess the correlation between the ordinal variables expressed by 1 to 10 Likert scale and the demographic variables. However, before performing the statistical test, all certain assumption for specific test was performed and none of them violated the statement. The results with p-values of less than 0.05 were counted statistically significant.

## Results

### Scio-demographic characteristics of live poultry/poultry-meat sellers

The socio-demographic characteristics of live poultry/poultry-meat sellers (n=432) in the study highlight several significant trends (Table 1). Notably, a substantial majority of participants were male (75.5%, p<0.01), age between 26 and 34 years (31.9%, p<0.01), majority having only primary (28.9%, p<0.01) education, had ≤5 years of experience (40.3%, p<0.01), with 43.3% earning less than 20,000 BDT monthly. Limited number of participations had training, 33.1% was trained on live poultry marketing (TLPM), and 28.5% had training on poultry meat processing and marketing (TPMPM).

**Table 1.**
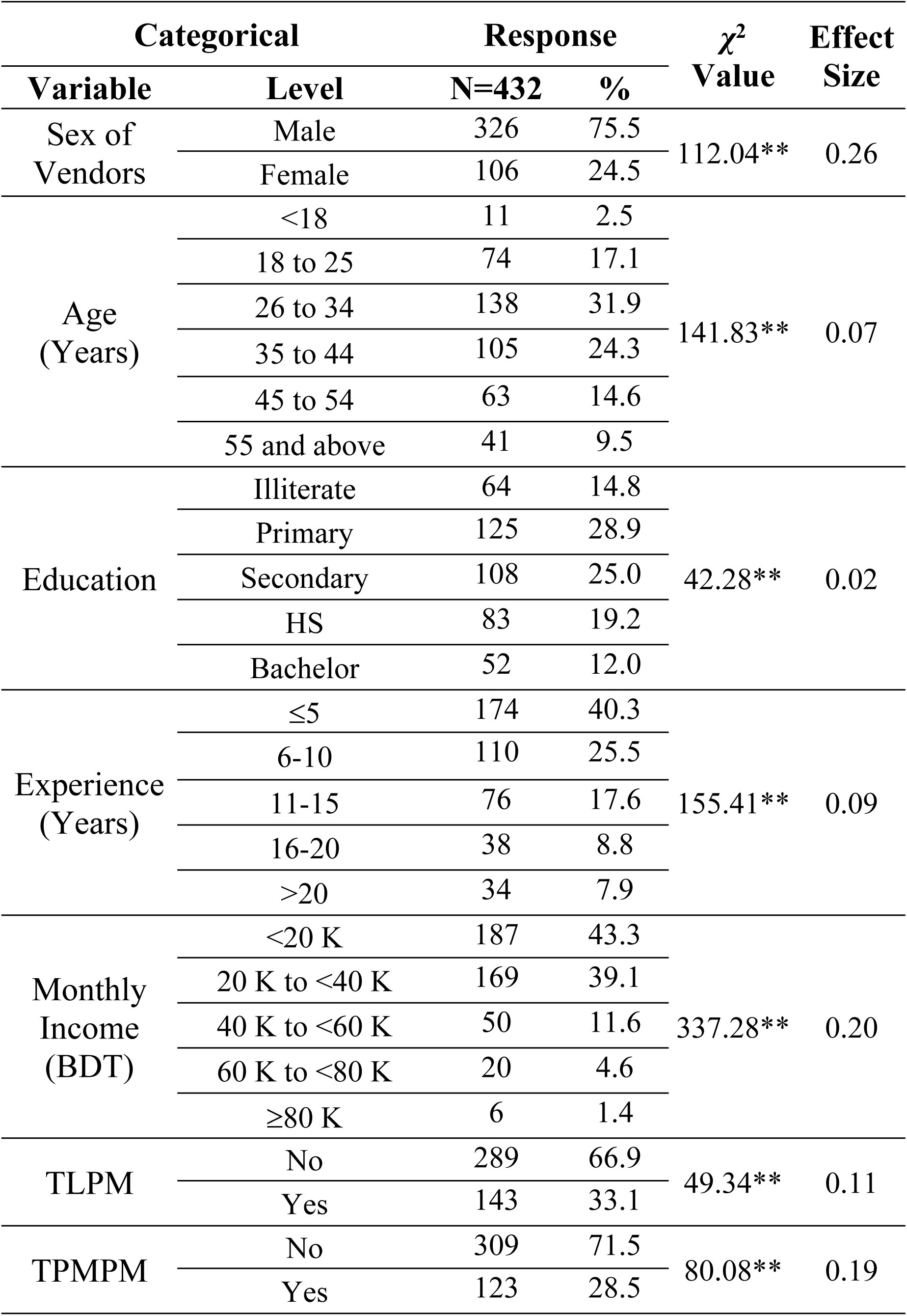
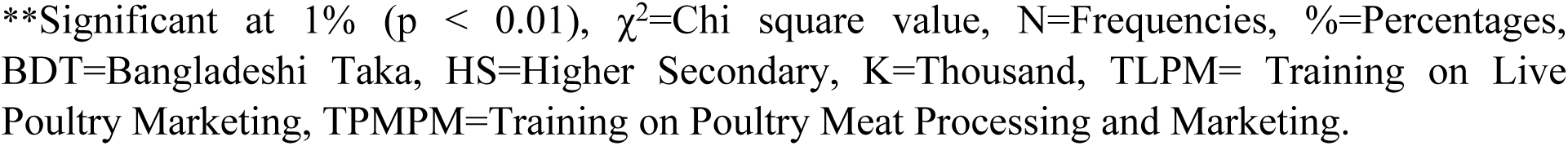
Scio-demographic characteristics of live poultry/poultry-meat sellers (n=432)

| Categorical | | Response | | $\chi^2$<br>Value | Effect<br>Size |
| --- | --- | --- | --- | --- | --- |
| Variable | Level | N=432 | % |  |  |
| Sex of<br>Vendors | Male | 326 | 75.5 | 112.04** | 0.26 |
|  | Female | 106 | 24.5 |  |  |
| Age<br>(Years) | <18 | 11 | 2.5 | 141.83** | 0.07 |
|  | 18 to 25 | 74 | 17.1 |  |  |
|  | 26 to 34 | 138 | 31.9 |  |  |
|  | 35 to 44 | 105 | 24.3 |  |  |
|  | 45 to 54 | 63 | 14.6 |  |  |
|  | 55 and above | 41 | 9.5 |  |  |
| Education | Illiterate | 64 | 14.8 | 42.28** | 0.02 |
|  | Primary | 125 | 28.9 |  |  |
|  | Secondary | 108 | 25.0 |  |  |
|  | HS | 83 | 19.2 |  |  |
|  | Bachelor | 52 | 12.0 |  |  |
| Experience<br>(Years) | ≤5 | 174 | 40.3 | 155.41** | 0.09 |
|  | 6-10 | 110 | 25.5 |  |  |
|  | 11-15 | 76 | 17.6 |  |  |
|  | 16-20 | 38 | 8.8 |  |  |
|  | >20 | 34 | 7.9 |  |  |
| Monthly<br>Income<br>(BDT) | <20 K | 187 | 43.3 | 337.28** | 0.20 |
|  | 20 K to <40 K | 169 | 39.1 |  |  |
|  | 40 K to <60 K | 50 | 11.6 |  |  |
|  | 60 K to <80 K | 20 | 4.6 |  |  |
|  | ≥80 K | 6 | 1.4 |  |  |
| TLPM | No | 289 | 66.9 | 49.34** | 0.11 |
|  | Yes | 143 | 33.1 |  |  |
| TPMPM | No | 309 | 71.5 | 80.08** | 0.19 |
|  | Yes | 123 | 28.5 |  |  |

\*\*Significant at 1% ( $p < 0.01$ ), $\chi^2$ =Chi square value, N=Frequencies, %=Percentages, BDT=Bangladeshi Taka, HS=Higher Secondary, K=Thousand, TLPM= Training on Live Poultry Marketing, TPMPM=Training on Poultry Meat Processing and Marketing.

### Evaluation of the knowledge response of the live poultry/poultry-meat sellers on risk of Zoonotic Salmonellosis

A significant proportion of participants (52.08%) were known about the topic of zoonotic diseases (Fig 2). Similarly, a significant (p<0.01) proportion (60.42%) positively responded on knowledge about Salmonellosis as a zoonotic disease with the highest effect size (ES) of 0.17. knowledge about food poisoning caused by zoonotic Salmonella were known to a significant proportion (58.56%) of poultry/poultry-meat sellers. Understanding of wild birds as a transmission route for zoonotic *Salmonella* (e.g., crows) was comparatively lower, with only 39.12% responding "Yes" (p<0.05) having a negligible effect size (0.01). Another critical area was knowledge about contamination risks through "dressed poultry meat" where 59.26% correctly identified with the effect size of 0.15. However, one of the most prominent results was the understanding of the risks impersonated by food and water intake in poultry shops, where 58.8% answered correctly (p<0.01; ES=0.15). A similar trend was observed for hand-washing practices before or during poultry dressing, where 55.32% (p<0.01, ES=0.11) acknowledged its role in reducing salmonellosis risk. The knowledge on the risks of using pond or river water to wash the dressed meat and utensils was 53.94% of poultry/poultry-meat sellers. Moreover, 52.08% had the knowledge on the washing utensils with detergent to eliminate *Salmonella spp.* Knowledge about the risks associated with repeated use of carcass dressing water showed a little lower proportion (49.77%, p<0.01, ES= 0.06) agreed that increase the risk of Salmonellosis. Proper disposal of poultry waste and byproducts as a preventive measure was significantly (p<0.01) identified by 56.02% of respondents with an effect size of 0.12.

**Fig 2.**
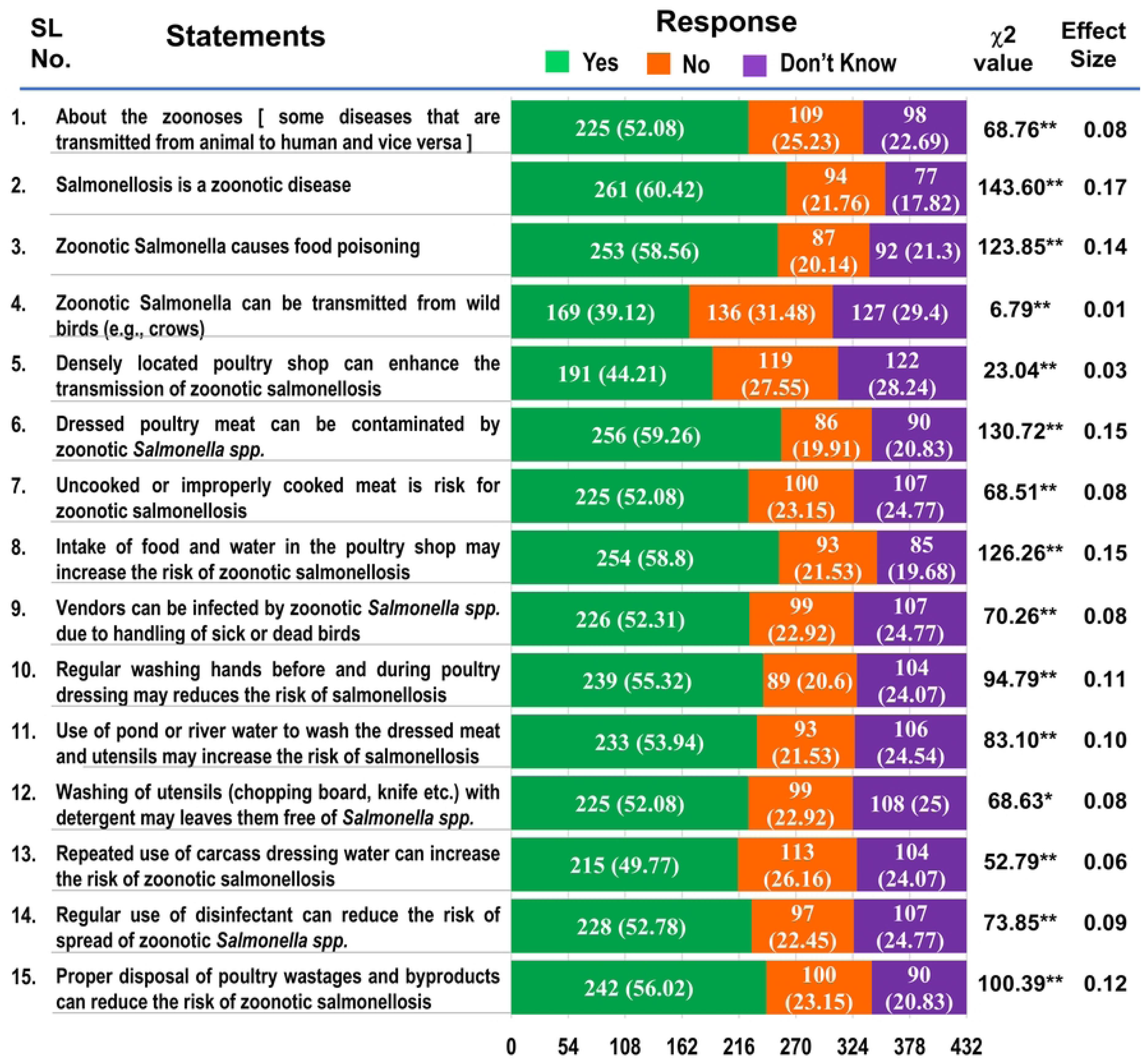
Evaluation of the knowledge response of the live poultry/poultry-meat sellers on risk of zoonotic Salmonellosis. ** Significant at 1% (p<0.01), * Significant at 1% (p<0.05), ***χ*^2^**= Chi square value, N=Frequencies, %= Percentages

### Association of demographic characteristics with the knowledge level of live poultry/poultry-meat sellers on the risk of zoonotic Salmonellosis

The association of demographic characteristics with the knowledge level of live poultry/poultry-meat sellers (n=432) revealed several significant findings, where 33.34% had the good knowledge and mean knowledge score is 53.12±0.89 (Table 2). In terms of gender, males demonstrated significantly (p<0.05) higher proportion of good knowledge (37.42%) compared to males (20.75). The mean knowledge score for males was significantly (p<0.05) higher at 54.5±1.03, compared to females (48.87±1.7). Individuals aged 55 years and above had significantly (p<0.01) highest proportion of good knowledge (43.9%). On the other hand, the age group of 26 to 34 years also showed significantly (p<0.01) highest mean knowledge score (57.01±1.57). Educational level was another strong determinant, individuals with a Bachelor’s degree (69.23%) and those with a higher secondary education (61.45%) exhibited significantly (p<0.01) better knowledge compared to illiterate individuals (18.75%) or those with only primary (21.60%) or secondary (16.67%) education. The mean knowledge scores were significantly (p<0.01) higher in the educated groups, reaching 67.69±1.98 for Bachelor’s degree holders and 65.94±2 for those with higher secondary education. Experience in poultry selling also influenced knowledge levels, indicating by the individuals having 16-20 years of experience showing the significantly (p<0.05) highest mean knowledge score (56.49±2.86). Additionally, TLPM and TPMPM had a significant (p<0.01) impact. Poultry sellers who received TLPM had higher mean knowledge score (59.49±1.62) compared to those without training (49.97±1.01). Similarly, TPMPM training was associated (p<0.01) with a higher knowledge score of 59.08±1.68, compared to those without training (50.74±1.01). Univariate and multivariate binary logistic regression analyses were conducted to explore factors associated with a good knowledge level of poultry/poultry-meat sellers (Table 2). In the univariate analysis, several variables were found to be significantly (p<0.01) associated with good knowledge, including gender (OR=0.44, 95% CI:0.26-0.74), age (OR=7.83, 95% CI:0.92-66.93 for individuals aged ≥55), educational level (OR=9.75, 95% CI:4.12-23.06 for Bachelor’s degree holders), TLPM (OR=3.2, 95% CI:2.10-4.89), and TPMPM (OR=2.43, 95% CI:1.58-3.75). In the multivariate analysis, the regression model was statistically significant (p<0.01), indicating that the model can differentiate between good and poor knowledge in considering the sellers demographic variable. Overall, this model could clarify between 22.3% (Cox and Snell R square) and 31.1% (Nagelkerke R squared) of the variance in poultry/poultry-meat sellers’ knowledge status. The model was also appropriately organized in 75.2% of instances. Hence, the goodness of fit for this model was determined by the p-value of 0.897 (p>0.05) in the “Hosmer and Lemeshow” test, which indicates that the final model is fit. In the results of logistic regression, after adjusting for other variables, age and educational level of the respondent remained a significant predictor of good knowledge. Poultry/poultry-meat sellers who are ≥55 years of age (OR=3.6, 95% CI:0.40-32.31) with a Bachelor’s degree (OR=6.64, 95% CI:2.59-17.04) ware significantly (p<0.01) more likely to have good knowledge. Similarly, those who received TLPM (OR=1.48, 95% CI:0.86-2.54) and TPMPM (OR=1.28, 95% CI:0.74-2.23) were more likely to have good knowledge, though this finding was marginally non-significant (p>0.05).

**Table 2.**
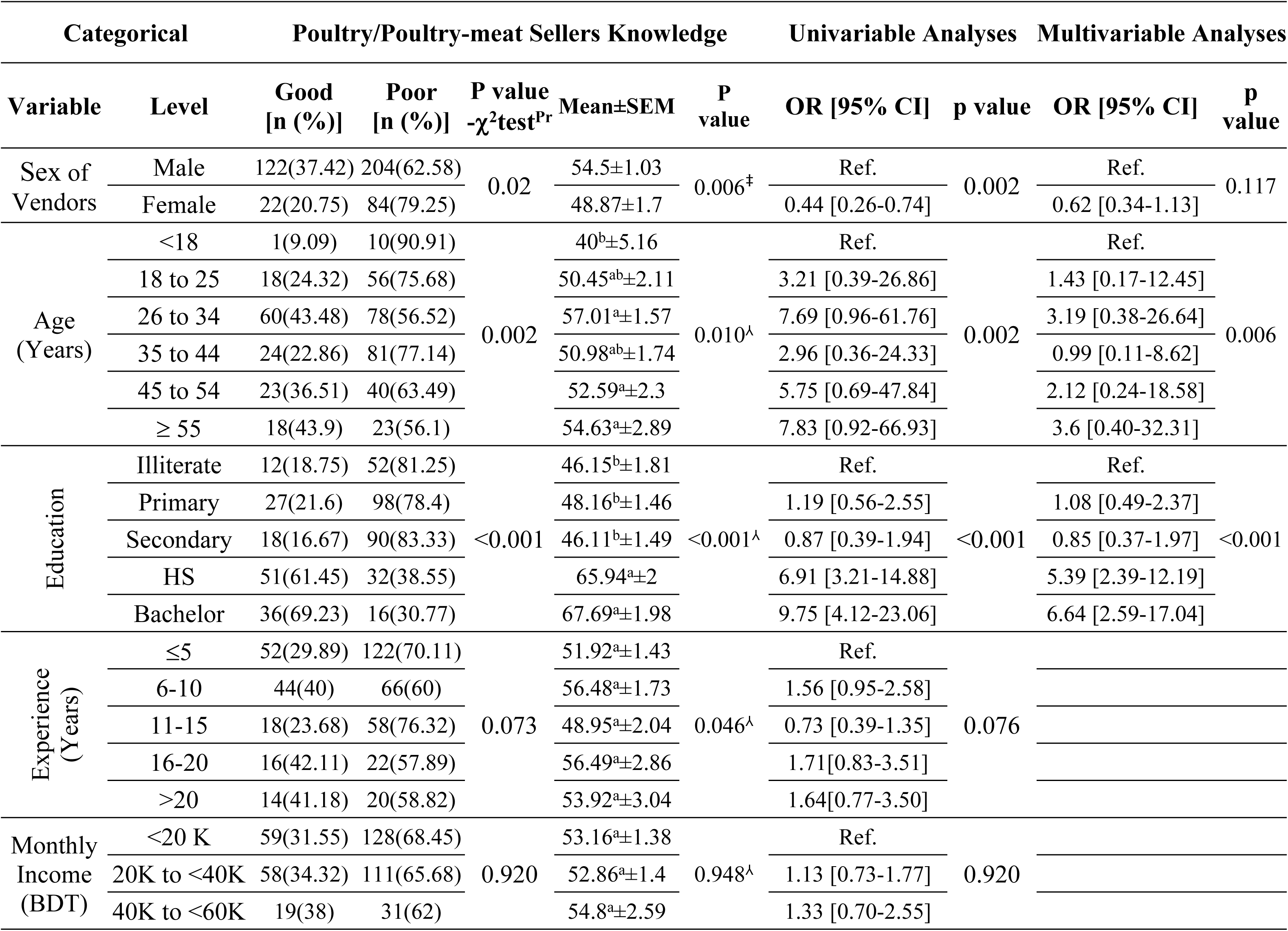
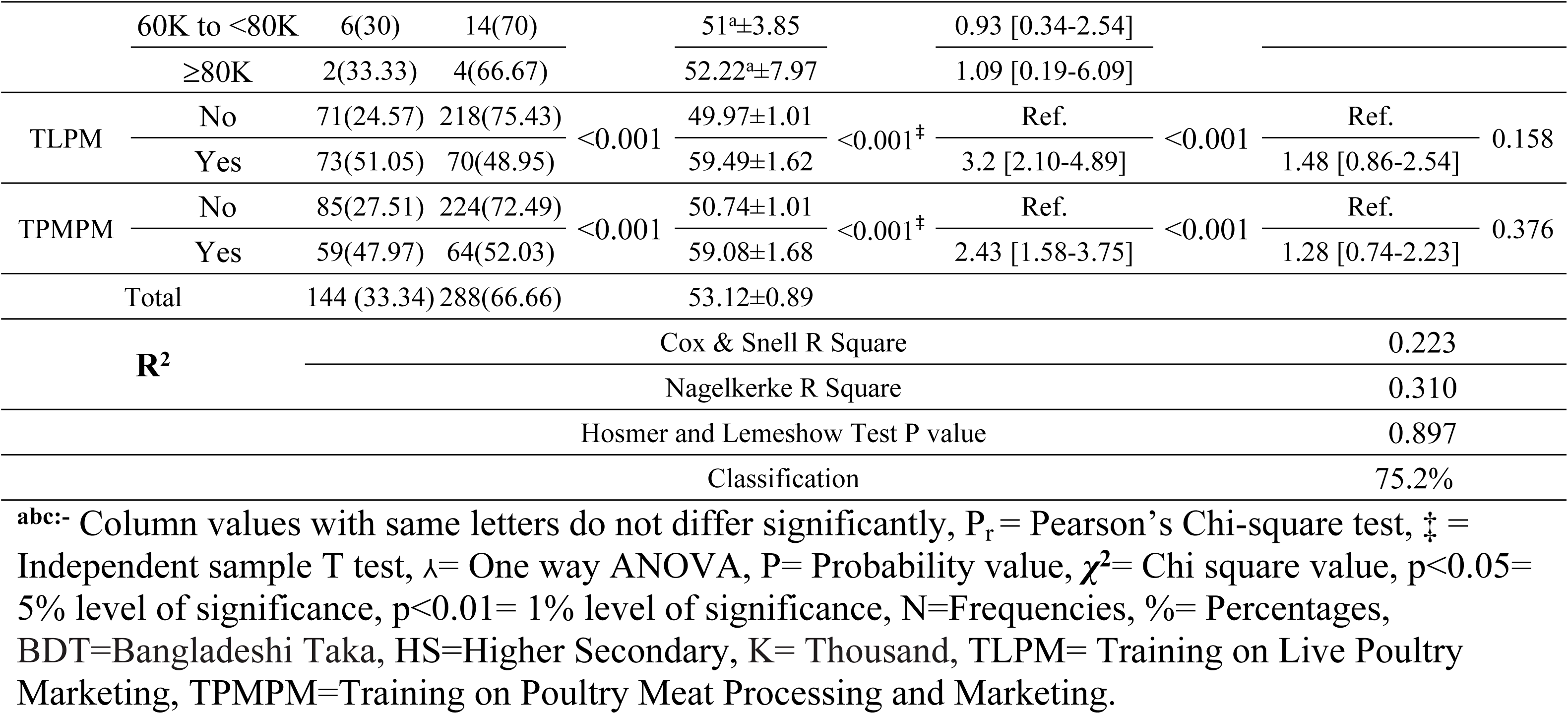
Association of demographic characteristics with the knowledge level of live poultry/poultry-meat sellers on the risk of zoonotic Salmonellosis and binary logistic regression analysis on their good knowledge score.

### Evaluation of the attitude’s response of the live poultry/poultry-meat sellers on risk of Zoonotic Salmonellosis

The attitudes responses of live poultry/raw poultry-meat sellers toward the risk of zoonotic salmonellosis are presented in the Fig 3. The calculated effect sizes (ranging between 0.05 to 0.16) validate the consistency of live poultry/poultry-meat sellers’ attitudes on risk of Zoonotic Salmonellosis. A significant (p<0.01, ES=0.13) proportion of respondents strongly agreed (42.59%) on training to reduce Salmonellosis. Similarly, significant (p<0.01, ES=0.09) proportion strongly agreed (34.26%), and agreed (24.54%) on good hygiene and sanitation to reduce the risk of Salmonella infection. Moreover, a significant (p<0.01, ES=0.09) proportion of participant (27.08%), agreed on “poultry meat should be dressed in a clean environment”, suggesting moderate acknowledgment of maintaining cleanliness. Nearly 28.7% strongly agreed, while 32.18% agreed to the use of clean safe water for dressing and washing of poultry meat demonstrated by a significant (p<0.01) effect size (ES=0.12). Approximately 34.26% of respondents strongly agreed, 28.24% agreed, and 16.44% moderately agreed on proper disposal of poultry offal’s (p<0.01, ES=0.11), emphasizing the sellers’ awareness of proper waste disposal as a biosecurity measure. A notable result (p<0.01, ES=0.16) was found that 42.59% of respondents strongly agreed and 25.23% agreed to workers should not rub hand on face, hair, etc. while dressing poultry meat. On the aspects of "Poultry meat dresser should not have long nails", 32.64% moderately agreed and 26.16% strongly agreed, yielding a significant effect (p<0.01, ES=0.11) of attitude toward the personal hygiene practices as critical to reducing zoonotic risks. The results significantly (p<0.01, ES=0.10) indicates that 30.32% strongly agreed, 28.94% moderately agreed, and 18.98% agreed on "Poultry meat dresser shouldn’t touch the money during processing of raw meat". This result highlights the sellers’ awareness of money handling as a contamination vector during meat processing. Proper cooking practices were widely recognized among sellers reflected by their significant (p<0.01, ES=0.09) response 24.54% strongly agreed and with 26.85% moderately agreed to poultry meat should be properly cooked.

**Fig 3.**
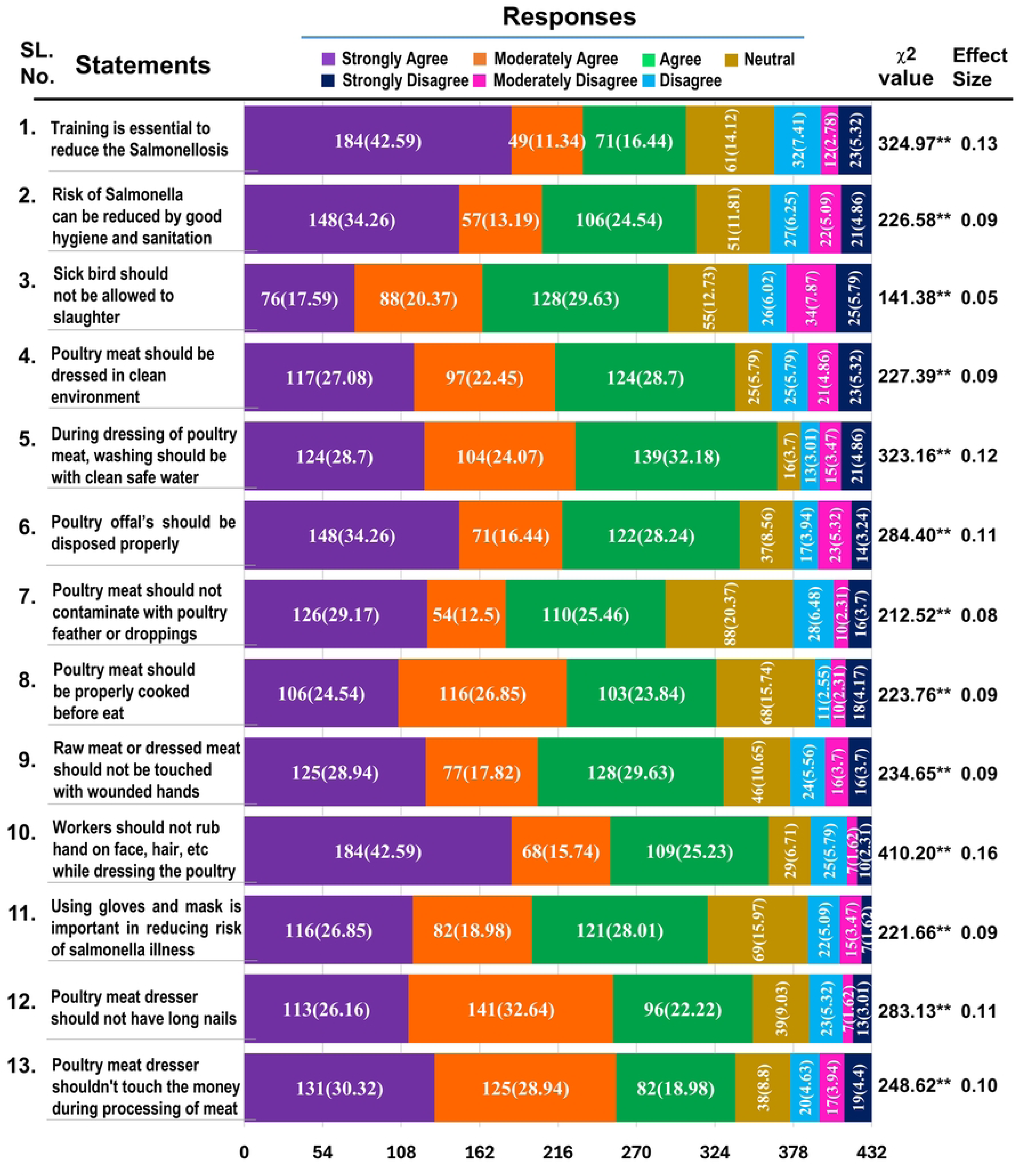
Evaluation of the Attitudes response of the live poultry/poultry-meat sellers on risk of zoonotic Salmonellosis. **Significant at 1% (p<0.01), *Significant at 1% (p<0.05), ***χ*^2^**= Chi square value, N=Frequencies, %= Percentages

### Association of demographic characteristics with the attitude level of live poultry/poultry-meat sellers on the risk of zoonotic Salmonellosis

The association between demographic characteristics and attitude levels (positive and negative) was statistically analysed, revealed that 62.03% of live poultry/poultry-meat sellers were bearing positive attitudes and mean attitude score was 68.59±0.79 (Table 3). Regarding gender, male sellers had a slightly higher proportion of positive attitudes (64.11%) compared to female sellers (55.66%), with a significant difference (p<0.05) in mean scores (69.52±0.91 for males, 65.72±1.51 for females). The sellers holding higher educational qualifications (HS and Bachelor degrees) showed significantly (p<0.01) higher positive attitudes. Those with a Bachelor’s degree had the highest proportion of positive attitudes (94.23%) and the highest mean score (78.48±1.47), which was notably (p<0.01) higher than the mean scores of illiterate sellers (63.06±1.83). Experience and monthly income did not show significant (p>0.05) associations with attitude levels. However, sellers who received TLPM were more likely to have positive attitudes (77.62%) compared to those who did not (54.33%), with a significant (p<0.01) mean score difference (72.99±1.22 for “yes” and 66.41±0.99 for “no” response). On the other hand, sellers with TPMPM also exhibited insignificantly (p>0.05) higher percentage of positive attitudes (68.29%). In the univariate binary logistic regression, education and TLPM emerged as significant (p<0.01) predictors of positive attitudes. Sellers with a Bachelor’s degree were over 16 times more likely to have a positive attitude (OR=16.33, 95% CI: 4.61-57.84), while HS educated sellers had over 6 times the odds (OR=6.55, 95% CI:2.94-14.59). Participants with TLPM had significantly (p<0.01) more tendency to positive attitude (OR=2.92, 95% CI:1.85-4.6). In multivariate analysis, participants with a Bachelor’s degree were where over 13 times more likely to have a positive attitude OR=13.48, 95% CI: 3.72-48.87). The regression model was significantly (p<0.01) differentiating between positive and negative attitudes with the clarification between 16.4% (Cox and Snell R square) and 22.3% (Nagelkerke R squared) of the variance in attitudes status. The model was also correctly arranged in 64.1% of instances. Hence, the final model was fit (“Hosmer and Lemeshow” test p value=0.83).

**Table 3.**
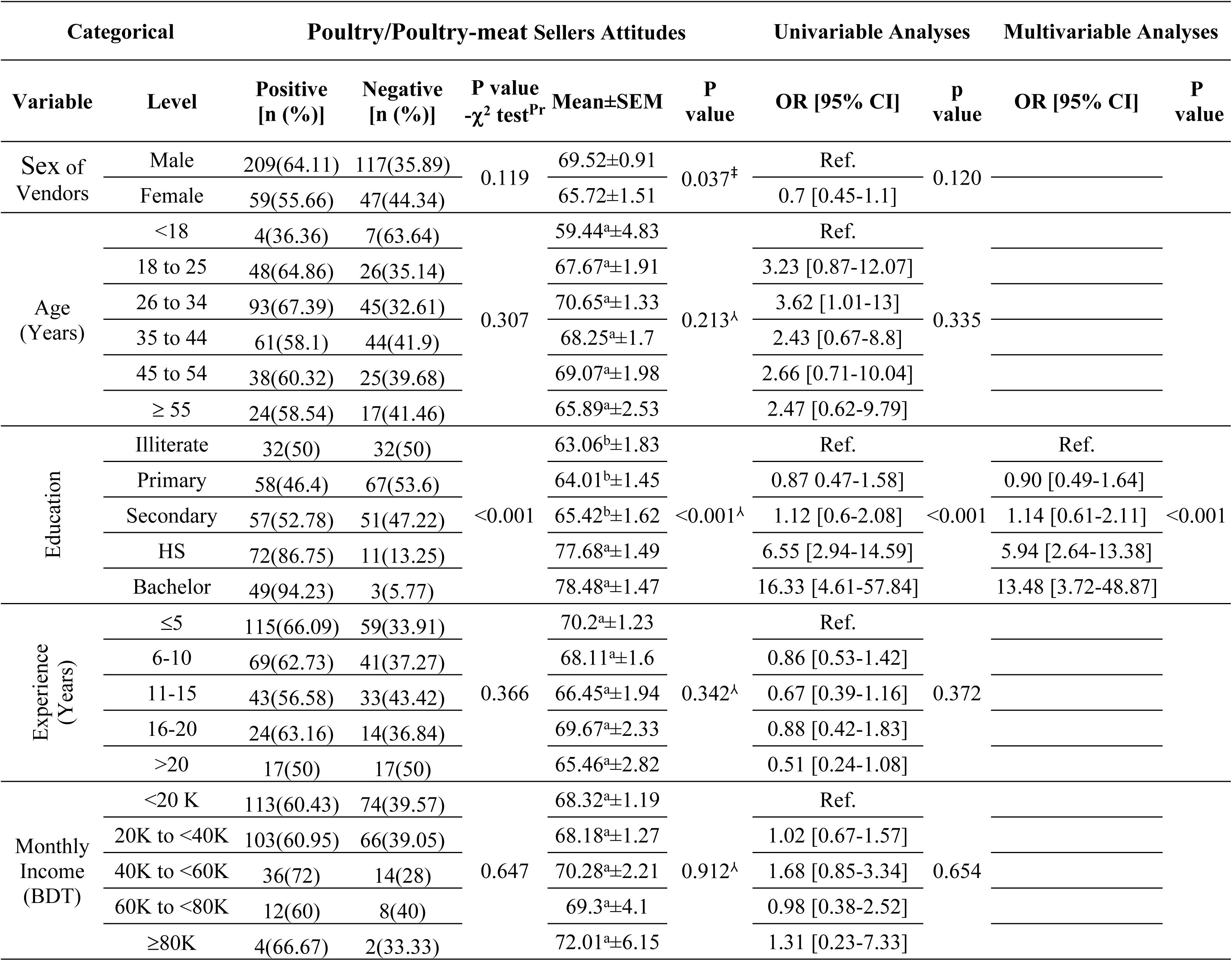
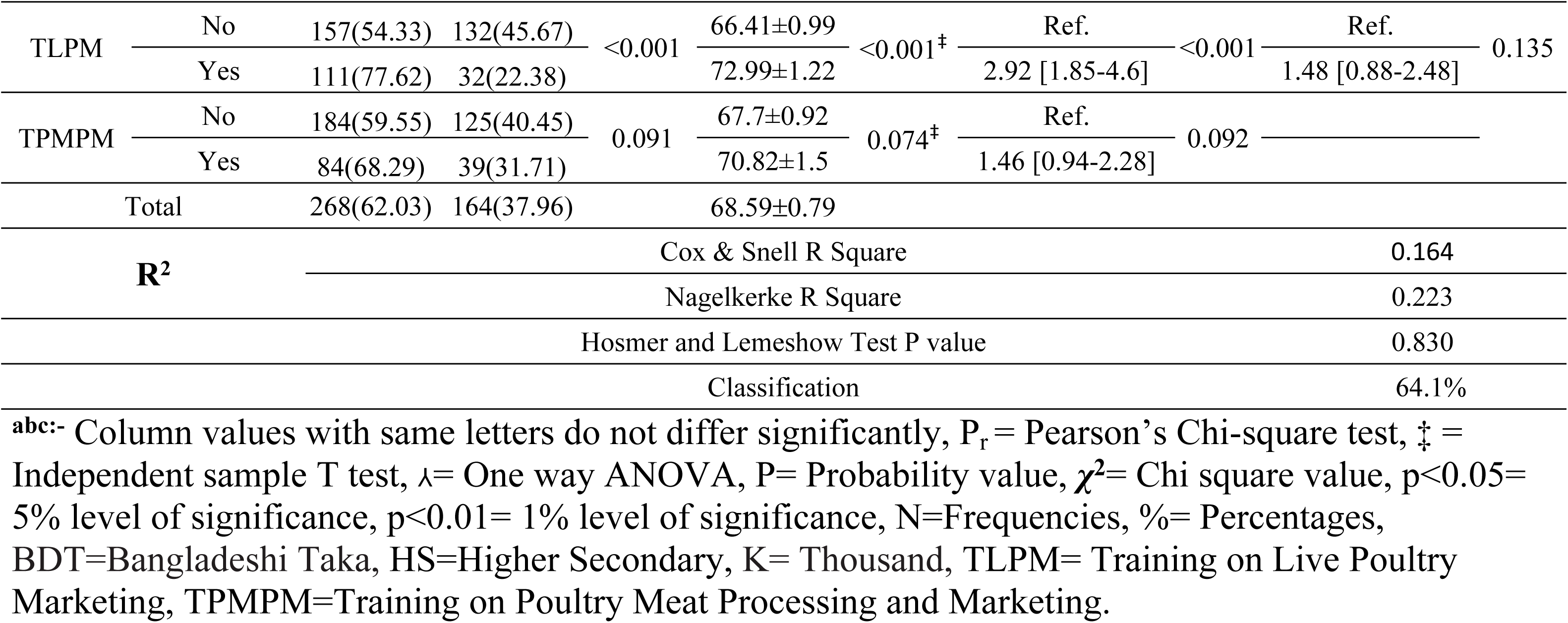
Association of demographic characteristics with the attitude level of live poultry/poultry-meat sellers on the risk of zoonotic Salmonellosis and binary logistic regression analysis on their positive attitude score.

### Evaluation of the biosecurity practice response of the live poultry/poultry-meat sellers on risk of Zoonotic Salmonellosis

The present study evaluates the biosecurity practices of poultry/poultry-meat sellers concerning the risk of zoonotic salmonellosis that is presented in Fig 4. Moderate to low effect sizes with high chi-square values across statements suggest practical significance of sellers’ responses with highlight areas where intervention, education, and strict monitoring are necessary to mitigating zoonotic risks. Higher proportion of the participants (36.34%, p<0.01, ES=0.03) reported that they always separated poultry according to age, type and health status (sickness) which is critical for disease containment. While 35.88% of respondents often refrained from taking food or drink in the shop, despite 19.91% never restricted that concerning a potential contamination risk (p<0.01, ES=0.02). Although 21.53% of respondents often restricted smoking in the shop, a large proportion (37.5%) did not comply with this practice (p<0.01, ES=0.04). With respect to hygiene practices, respondent reported they always (14.81%) and often (21.53%) using fresh water during dressing and cleaning (p<0.01, ES=0.03), while 36.57% always used hot water (p<0.01, E=0.06) to minimize the microbial contamination. Hand hygiene was well-practiced by 38.19% (p<0.01, ES=0.05) who every time washed hands with soap before and after processing meat. Regarding structural cleanliness and utensil hygiene, 33.1% of participants often cleaned and washed tools like knives and chopping boards before and after use (p<0.01, ES=0.02), and 36.34% usually cleaned the shop with disinfectants at closing time (p<0.01, ES=0.06). Avoiding Jewellery, often practiced in poultry/poultry-meat shop, was reported by 39.58% of sellers (p<0.01, ES=0.06). Use of personal protective equipment (PPE) was less prevalent. Only 15.74%, 24.54% and 34.03% of sellers reported always, often, and rarely using masks/aprons/caps respectively (p<0.01, ES=0.02). Additionally, only 14.58% reported that they use hand gloves always. However, changing of clothes after meat processing was more common, poultry/poultry-meat sellers always practiced by 36.11% (p<0.01, ES=0.04). Proper waste disposal also showed encouraging trends. A total of 35.88% often disposed the slaughtering waste hygienically where 24.31% dispose regularly (p<0.01, ES=0.03). The higher proportion of sellers (40.28%) always used specific dustbins for blood, viscera, and feathers of poultry carcass (p<0.01, ES=0.06). Similarly, 37.5% often followed hygienic methods for dead bird disposal where 23.15% practiced regularly (p<0.01, ES=0.03). Vector control and access restriction practices were also addressed. About 37.5% always employed measures to control flies and rats (p<0.01, ES=0.06), while 36.57% often restricted wild birds in the slaughtering area (p<0.01, ES=0.06). However, efforts to prevent the presence of dogs and cats near the shop were less consistent, with only 24.07% regularly and 28.94% rarely practicing such restrictions (p<0.01, ES=0.01).

**Fig 4.**
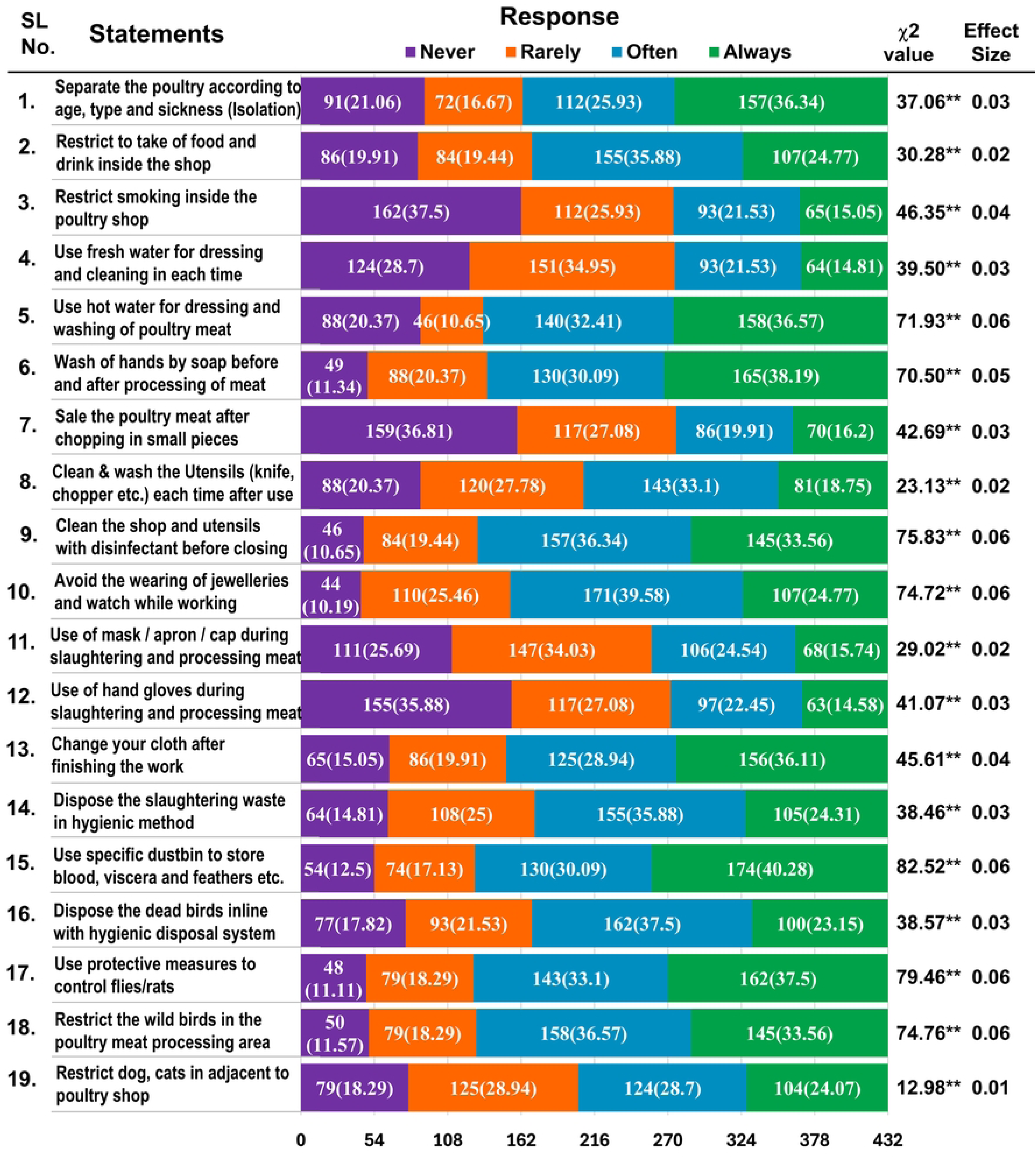
Evaluation of the biosecurity practice response of the live poultry/poultry-meat sellers on risk of zoonotic Salmonellosis. **Significant at 1% (p<0.01), *Significant at 1% (p<0.05), *χ*^2^= Chi square value, N=Frequencies, %= Percentages

### Association of demographic characteristics with the biosecurity practice level of live poultry/poultry-meat sellers on the risk of zoonotic Salmonellosis

The association of demographic characteristics with biosecurity practices among poultry and poultry meat sellers are presented in the Table 4. A total of 41.43% of poultry/poultry-meat sellers were involved in correct practice and the mean of biosecurity practices was 54.58±0.73. Regarding gender, male sellers (46.01%) had significantly (p<0.01) higher proportion of correct biosecurity practices compared to female sellers (27.36%) with a significant difference (p<0.01) in mean scores (55.94±0.85 for males and 50.4±1.37 for females). The highest significant (p<0.05) mean practice (57.18±1.28) score was observed among the 26 to 34 yeas ages sellers. The sellers holding HS showed significantly (p<0.01) more correct biosecurity practices (69.88%) with the highest mean score (65.8±1.31). Experience and monthly income did not show significant associations with correct biosecurity practices levels. However, sellers who received TLPM were significantly (p<0.01) higher in correct biosecurity practices (55.24%) compared to those who did not (34.6%), with a significant (p<0.01) mean score difference (59.91±1.24 for yes and 51.95±0.86 for no response). Similarly, respondent with TPMPM was significantly (p<0.05) higher proportion of correct biosecurity practices (50.41%) than without TPMPM (37.86%). The mean score was also significantly (p<0.01) higher for having TPMPM (59.06±1.29) than no TPMPM (52.8±0.86). In the univariate analysis, several variables were found to be significantly (p<0.01) associated with correct biosecurity practices, including gender (OR=0.44, 95% CI:0.27-0.71 for female), educational level (OR=5.75, 95% CI:2.58-12.83 for Bachelor’s degree and OR=5.93, 95% CI: 2.89-12.17 for HS holders), TLPM (OR=2.33, 95% CI:1.55-3.51), and TPMPM (OR=1.67, 95% CI:1.09-2.54). Age, experience and income were not significantly associated with biosecurity practices. In multivariate analysis, female vendors were less likely to follow correct biosecurity practices (OR=0.54, 95% CI:0.32–0.91, p<0.05). Education level, HS (OR=5.52, 95% CI:2.62–11.63) and bachelor’s education (OR=4.81, 95% CI:2.03– 11.39) remained significant (p<0.01) predictors of correct biosecurity practices. The effect of TLPM (OR=1.27, 95% CI:0.76–2.11) and TPMPM (OR=0.80, 95% CI:0.48–1.35) was not significant (p>0.05) in multivariate analysis though these were significant in the univariate analysis. However, the multivariate regression model was significantly (p<0.01) differentiating between correct and incorrect practices with the clarification between 15.4% (Cox and Snell R square) and 20.7% (Nagelkerke R squared) of the variance in biosecurity practices. The model was also correctly arranged in 70.8% of instances. Hence, the final model was fit (“Hosmer and Lemeshow” test p value=0.268).

**Table 4.** Association of demographic characteristics with the biosecurity practice level of live poultry/poultry-meat sellers on the risk of zoonotic Salmonellosis and binary logistic regression analysis on their correct biosecurity practices score.

| Categorical |  | Biosecurity Practices by Poultry/Poultry-meat Sellers |  |  |  | Univariable Analyses |  | Multivariable Analyses |  |  |
| --- | --- | --- | --- | --- | --- | --- | --- | --- | --- | --- |
| Variable | Level | Correct<br>[n (%)] | Incorrect<br>[n (%)] | P value<br>$\chi^2$ test<sup>Pr</sup> | Mean±SEM | P<br>value | OR [95% CI] | P<br>value | OR [95% CI] | P<br>value |
| Sex of Vendors | Male | 150(46.01) | 176(53.99) | <0.001 | 55.94±0.85 | 0.001<sup>‡</sup> | Ref. | <0.001 | Ref. | 0.02 |
|  | Female | 29(27.36) | 77(72.64) |  | 50.4±1.37 |  | 0.44 [0.27-0.71] |  | 0.54 [0.32-0.91] |  |
| Age (Years) | <18 | 2(18.18) | 9(81.82) | 0.125 | 45.93<sup>b</sup>±4.72 | 0.045<sup>λ</sup> | Ref. | 0.140 |  |  |
|  | 18 to 25 | 25(33.78) | 49(66.22) |  | 51.75<sup>ab</sup>±1.84 |  | 2.30 [0.46-11.44] |  |  |  |
|  | 26 to 34 | 67(48.55) | 71(51.45) |  | 57.18<sup>ab</sup>±1.28 |  | 4.25 [0.89-20.37] |  |  |  |
|  | 35 to 44 | 47(44.76) | 58(55.24) |  | 55.36<sup>ab</sup>±1.38 |  | 3.65 [0.75-17.70] |  |  |  |
|  | 45 to 54 | 23(36.51) | 40(63.49) |  | 52.99<sup>ab</sup>±1.88 |  | 2.59 [0.51-13.02] |  |  |  |
|  | ≥ 55 | 15(36.59) | 26(63.41) |  | 53.7<sup>a</sup>±2.48 |  | 2.60 [0.49-13.64] |  |  |  |
| Education | Illiterate | 18(28.13) | 46(71.88) | <0.001 | 48.41<sup>b</sup>±1.89 | <0.001<sup>λ</sup> | Ref. | <0.001 | Ref. | <0.001 |
|  | Primary | 33(26.4) | 92(73.6) |  | 48.77<sup>b</sup>±1.19 |  | 0.92 [0.47-1.8] |  | 0.88 [0.44-1.74] |  |
|  | Secondary | 34(31.48) | 74(68.52) |  | 51.09<sup>b</sup>±1.32 |  | 1.17 [0.60-2.32] |  | 1.20 [0.60-2.38] |  |
|  | HS | 58(69.88) | 25(30.12) |  | 65.8<sup>a</sup>±1.31 |  | 5.93 [2.89-12.17] |  | 5.52 [2.62-11.63] |  |
|  | Bachelor | 36(69.23) | 16(30.77) |  | 65.49<sup>a</sup>±1.58 |  | 5.75 [2.58-12.83] |  | 4.81 [2.03-11.39] |  |
| Experience (Years) | ≤5 | 67(38.51) | 107(61.49) | 0.433 | 53.86<sup>a</sup>±1.18 | 0.588<sup>λ</sup> | Ref. | 0.435 |  |  |
|  | 6-10 | 51(46.36) | 59(53.64) |  | 56.46<sup>a</sup>±1.33 |  | 1.38 [0.85-2.24] |  |  |  |
|  | 11-15 | 27(35.53) | 49(64.47) |  | 53.74<sup>a</sup>±1.58 |  | 0.88 [0.50-1.54] |  |  |  |
|  | 16-20 | 18(47.37) | 20(52.63) |  | 53.14<sup>a</sup>±2.92 |  | 1.44 [0.71-2.91] |  |  |  |
|  | >20 | 16(47.06) | 18(52.94) |  | 55.68<sup>a</sup>±2.87 |  | 1.42 [0.68-2.97] |  |  |  |
| Monthly Income (BDT) | <20 K | 75(40.11) | 112(59.89) | 0.702 | 53.69<sup>a</sup>±1.09 | 0.637<sup>λ</sup> | Ref. | 0.737 |  |  |
|  | 20K to <40K | 69(40.83) | 100(59.17) |  | 55<sup>a</sup>±1.15 |  | 1.03 [0.67-1.57] |  |  |  |
|  | 40K to <60K | 23(46) | 27(54) |  | 56.18<sup>a</sup>±2.14 |  | 1.272 [0.68-2.39] |  |  |  |
|  | 60K to <80K | 8(40) | 12(60) |  | 53.42<sup>a</sup>±4.09 |  | 1.00 [0.39-2.55] |  |  |  |
|  | ≥80K | 4(66.67) | 2(33.33) |  | 61.11<sup>a</sup>±8.98 |  | 2.99 [0.53-16.72] |  |  |  |
| TLPM | No | 100(34.6) | 189(65.4) | <0.001 | 51.95±0.86 | <0.001<sup>‡</sup> | Ref. | <0.001 | Ref. | 0.356 |
|  | Yes | 79(55.24) | 64(44.76) |  | 59.91±1.24 |  | 2.33 [1.55-3.51] |  | 1.27 [0.76-2.11] |  |
| TPMPM | No | 117(37.86) | 192(62.14) | 0.017 | 52.8±0.86 | <0.001<sup>‡</sup> | Ref. | 0.017 | Ref. | 0.406 |
|  | Yes | 62(50.41) | 61(49.59) |  | 59.06±1.29 |  | 1.67 [1.09-2.54] |  | 0.80 [0.48-1.35] |  |
| Total |  | 179 (41.43) | 253 (58.56) |  | 54.58±0.73 |  |  |  |  |  |
| R<sup>2</sup> |  | Cox & Snell R Square |  |  |  |  |  |  | 0.154 |  |
|  |  | Nagelkerke R Square |  |  |  |  |  |  | 0.207 |  |
|  |  | Hosmer and Lemeshow Test P value |  |  |  |  |  |  | 0.268 |  |
|  |  | Classification |  |  |  |  |  |  | 70.8% |  |

<sup>ab</sup>:- Column values with same letters do not differ significantly, P<sub>r</sub> = Pearson's Chi-square test, ‡ = Independent sample T test, λ = One way ANOVA, P = Probability value, χ<sup>2</sup> = Chi square value, p<0.05 = 5% level of significance, p<0.01 = 1% level of significance, N = Frequencies, % = Percentages, BDT = Bangladeshi Taka, HS = Higher Secondary, K = Thousand, TLPM = Training on Live Poultry Marketing, TPMPM = Training on Poultry Meat Processing and Marketing.

### Scatter diagrams showing the Pearson linear correlations of knowledge, attitudes, and practices score of live poultry/poultry-meat seller

The Pearson linear correlation analysis reveals significant (p<0.01) associations between knowledge Score, attitudes score, and practices score of live poultry/raw poultry-meat sellers (n=432), as indicated in the scatter diagrams (Fig 5). Panel A demonstrates a significant (p<0.01) low-strength of positive correlation between knowledge score and attitudes score with a linear regression equation (y=58.16+0.2*x) and correlation (r=0.221). Similarly, Panel B showed a significant (p<0.01) low-strength of positive correlation between Knowledge Score and Practices Score, described by the equation y=42.3+0.23*x, with a correlation, r=0.281. The strongest association is observed in Panel C, where attitudes score had moderate-strength of positive correlation with practices score. The regression equation was y=46.79+0.4*x, with r=0.371 and p<0.01, reflecting a stronger linear relationship.

**Fig 5.**
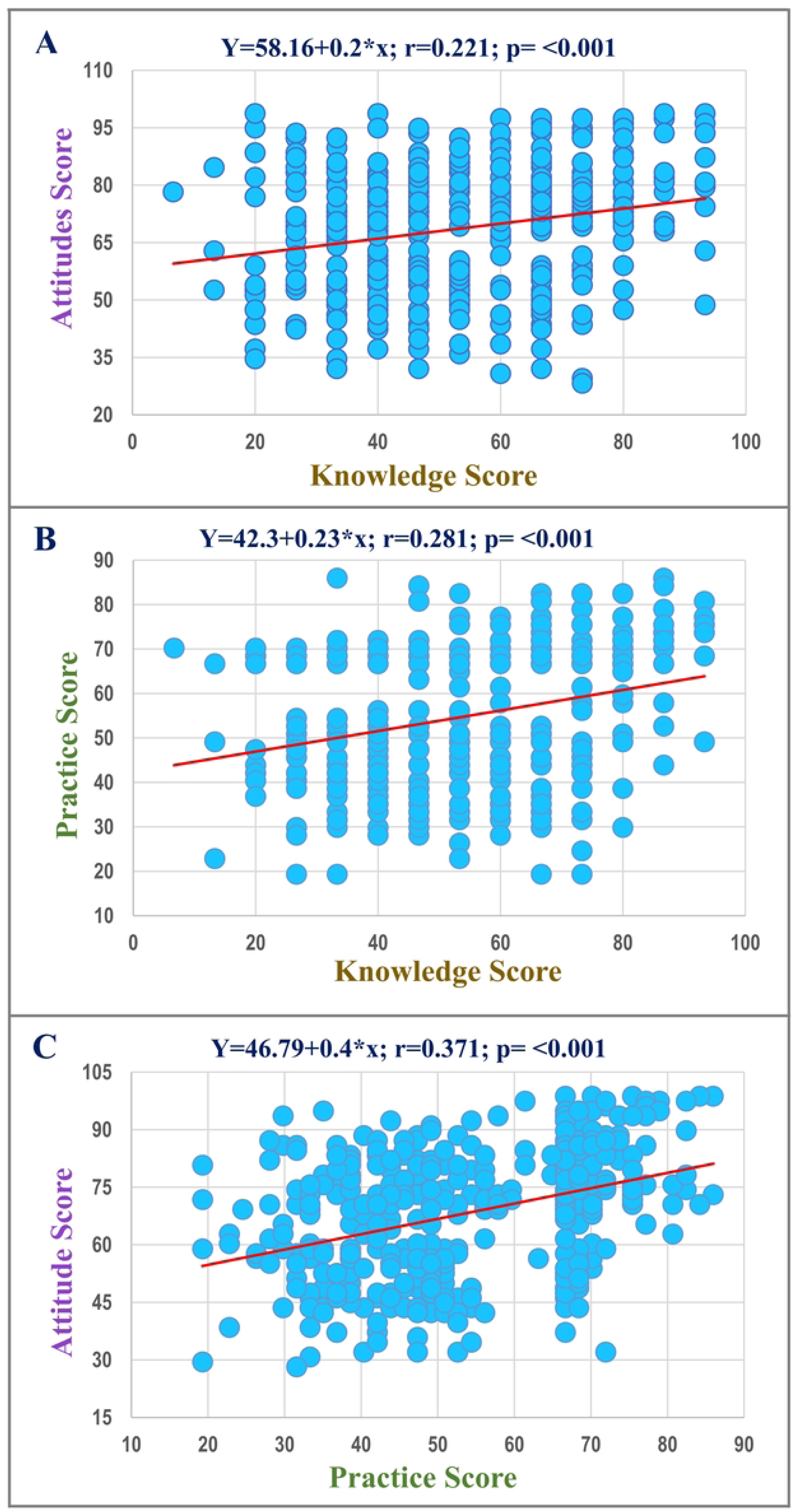
Scatter diagrams showing the Pearson linear correlations of (A) knowledge score with attitudes score, (B) knowledge score with practices score, and (C) attitudes score with practices score of live poultry/poultry-meat seller. p<0.05= 5% level of significance, p<0.01= 1% level of significance

### Perceived Risk, Biosecurity Confidence, and Education Interest by KAP and Demographics

The box plot showing the Likert scale (1=very low, 10=very high) of participant’s perceived risk of zoonotic disease transmission (PPRZDT), participant’s confidence in personal biosecurity measures (PCPBM) and participant’s interest on zoonoses education and training (PIZETP) for prevention of live poultry/poultry-meat seller on zoonotic salmonellosis according to (A) knowledge, (B) attitude, and (C) biosecurity practices (Fig 6). The poultry/poultry-meat seller who had good knowledge, scored themselves to have significantly (p<0.01) more perception on risk of zoonotic disease transmission where the median was 7 (Fig 6A). Similarly, poultry/poultry-meat seller with positive attitude and involved in correct biosecurity practice showed significantly (p<0.05) more perception on risk of zoonotic disease transmission indicated by the median of 7 (Fig 6B and Fig 6C). The small effect sizes for confidence and interest further support the lack of meaningful differences between these groups. The correlation analysis with demographic variables (Fig 6D) revealed several notable associations. PPRZDT was significantly correlated with education (r=0.131, p<0.01) and TLPM (r=0.084, p<0.05). PCPBM showed a significant positive correlation with gender (r=0.108, p<0.01), indicating higher confidence among certain gender groups. PIZETP demonstrated no significant correlations with demographic variables. These results underscore the role of education, income, and gender in influencing perceptions and attitudes toward zoonotic disease prevention.

**Fig 6.**
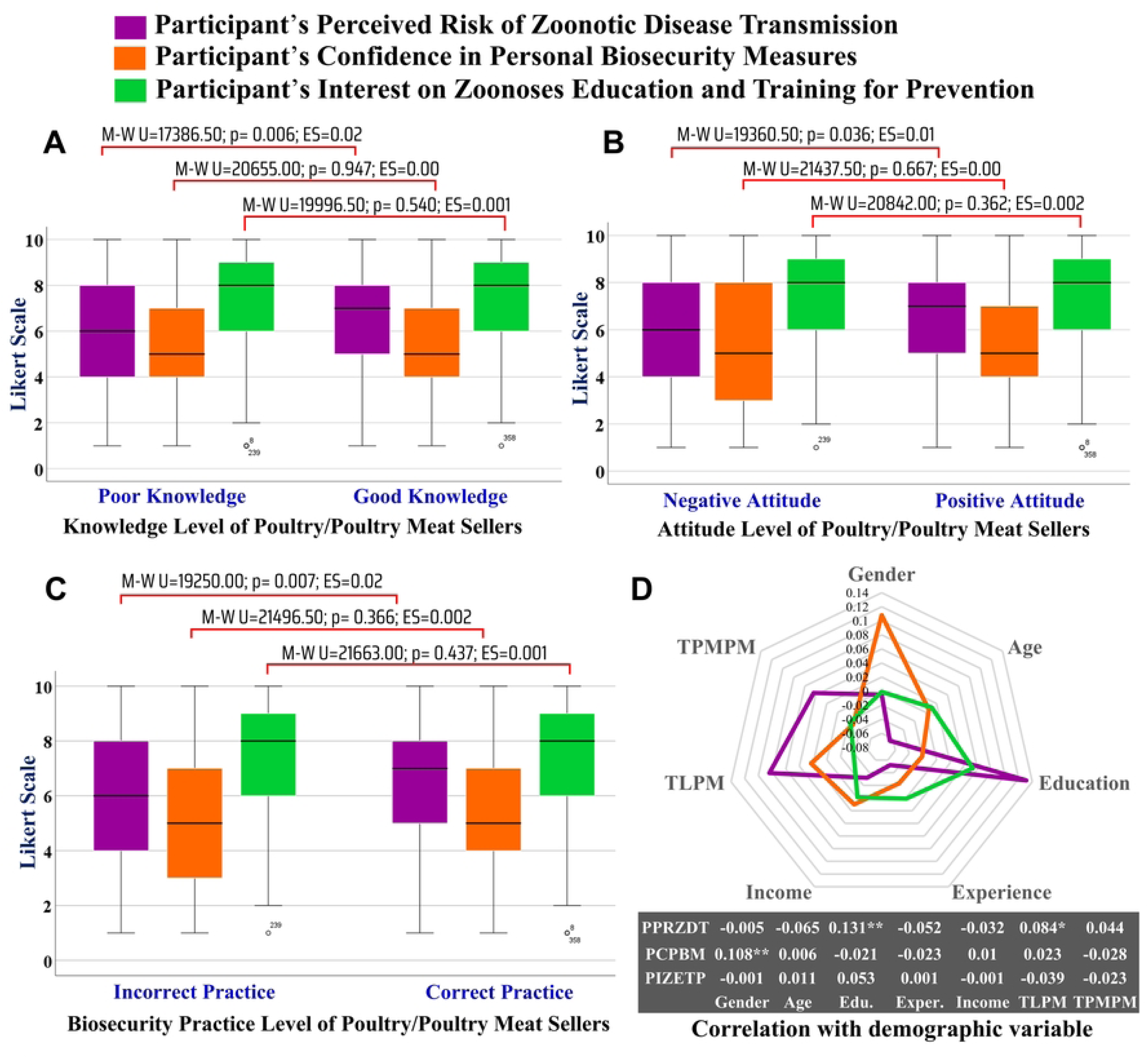
The box plot showing the Likert scale of perceived risk of zoonotic disease transmission, confidence in personal biosecurity measures, and interest on zoonoses education and training for prevention of live poultry/poultry-meat seller on zoonotic salmonellosis according to (A) knowledge, (B)attitude, (C) biosecurity practices of live poultry/poultry-meat seller, and (D) the Kendall’s tau-b correlation of Likert scale variable with the demographic variable. Highly significant at 1% (p *<* 0.01), Significant at 5% (p *<* 0.05), p = Probability value, M-W U= Value of Mann–Whitney test.

## Discussion

The present study investigates live poultry/poultry-meat seller’s perception, attitudes and biosecurity practices to mitigate the risk of salmonellosis. In every year’s considerable portion of live poultry vending shops are increasing in Bangladesh, has created a new field of study to understand the risk of more prevalent zoonotic disease, such as salmonellosis transmission [12,35]. However, a lack of sufficient studies has made it unknown how poultry/poultry-meat sellers are managing this risk issue. This study highlighted the notable socio-demographic patterns which has influence on biosecurity practices, perception, and awareness of live poultry/poultry-meat sellers to mitigate the risk of zoonotic salmonellosis. To facilitate interpretation of the study’s outcomes, Fig. 7 presents a graphical synthesis of the research design and major findings related to the knowledge, attitudes, and practices of live poultry/poultry-meat sellers. The majority of sellers in this study were male, young (aged 26–34 years), with only primary education and limited experience in poultry trading. These findings align with previous studies in similar urban and peri-urban settings, where poultry trade is predominantly male-driven, informal, and often not supported by structured education or training opportunities [14,36,37]. Actually, negligible number of females are involved in poultry selling particularly they are in the less busy markets. However, over half (60.42%) of the total respondents recognized salmonellosis as a zoonotic disease. Some earlier studies reported that about 94% of participants from Ireland understood that Salmonella could cause illness in both poultry and humans [38]. In another study reported that the most respondents from Nigeria had limited awareness of Salmonella infections and their effects on poultry [39]. The relatively higher level of awareness observed in our study may be attributed to public health campaigns and increased digital information access. Nonetheless, a higher proportion of participants had a general understanding of Salmonella’s impact, but there remains a substantial gap in understanding its zoonotic potential, modes of transmission, and preventive measures [40]. Although more than half of the respondents acknowledged food poisoning risks from zoonotic Salmonella and recognized the contamination risk of dressed poultry meat. These findings have little bit difference from other study, were reported that only 28.3 % of meat handlers in Gopalganj of Bangladesh had adequate food safety knowledge [23]. These dissimilarities might be due to regional differences inside a country. In our study, the low knowledge regarding wild birds as vectors may be due to the limited inclusion of environmental transmission routes in informal training or day-to-day experience, as sellers often focus on direct poultry handling rather than indirect sources of contamination. Although a majority recognized the importance of hand washing during meat processing (55.32%) and risks of water contamination (53.94%), a significant portion remained unaware of the dangers associated with reusing carcass dressing water or using pond/river water for cleaning. These results underscore the persistent challenges in translating general health awareness into context-specific biosecurity knowledge, as noted in previous studies [14,23,27,28]. Particularly, the hhygienic knowledge depend on the educational status and the experiences of the vendors [41]. The overall knowledge score of vendors in our study underscores the need for increased educational programs, and training initiatives among the sellers, as highlighted in previous study [32]. Participants of older ages, and those holding a Bachelor’s degree, were more likely to possess good knowledge. These associations are consistent with previous research showing that education enhances awareness and interpretation of zoonotic risks [14]. In another study conducted in Bangladesh reported that older meat sellers exhibiting superior knowledge on meat hygiene compared to the younger [27], which support our findings. Logistic regression analyses further reinforced the role of education and age as significant predictors of knowledge. Sellers who attended training on live poultry marketing (TLPM) and poultry meat processing and marketing (TPMPM) scored significantly higher in knowledge. Similar effects of training have been observed among meat sellers on other study [27], where structured training programs improved knowledge of meat handlers on meat hygiene enhancing their safety practices particularly newcomers with limited knowledge. Although training (TLPM and TPMPM) did not remain significant in the multivariate model, the effect sizes suggest a trend toward improved knowledge with training exposure. This indicates that while formal education plays a foundational role, targeted training can serve as a valuable supplement. Perhaps, the frequent professional training could help to improve knowledge among meat sellers, mitigating zoonotic transmission risks and ensuring biosecurity practices [25].

**Fig 7.**
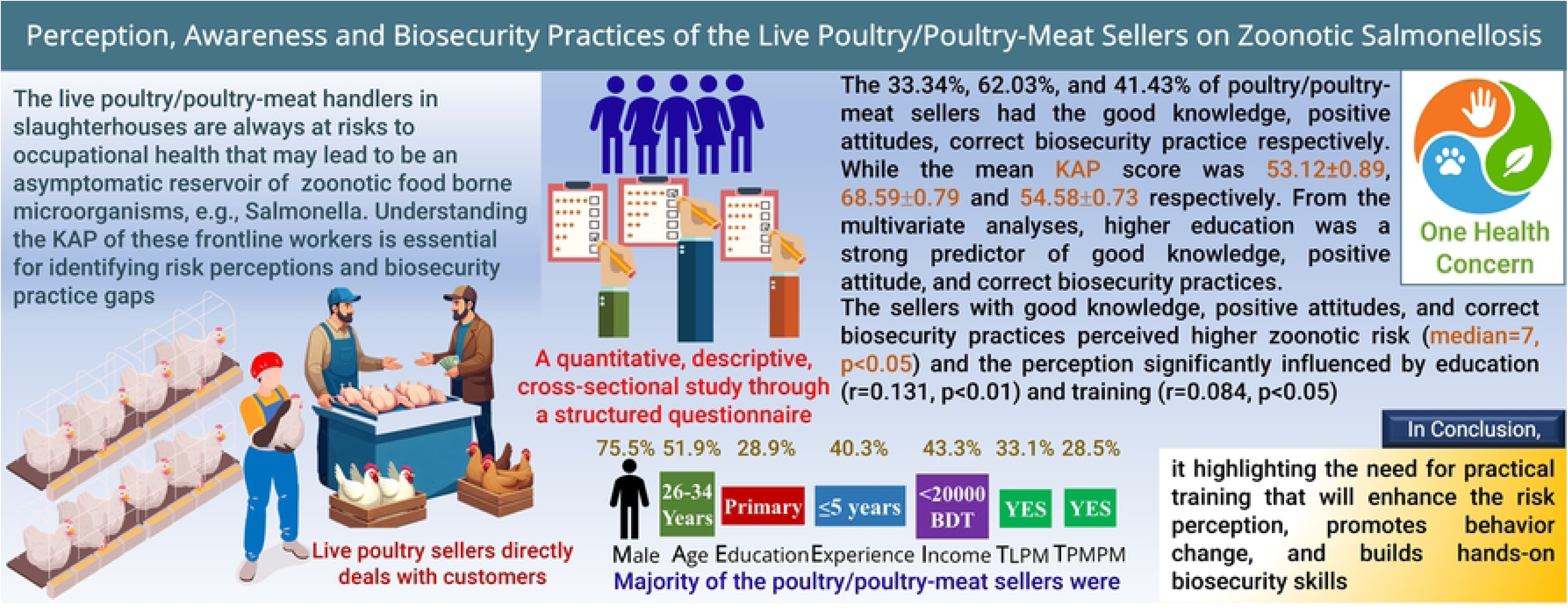
Graphical Illustration of live poultry/poultry-meat seller’s perception, awareness and biosecurity practices to mitigate the risk of zoonotic salmonellosis.

The present study revealed that live poultry and raw poultry-meat sellers generally hold moderately positive attitudes toward the prevention of zoonotic salmonellosis. The consistency of attitude responses suggests that these sellers possess a basic awareness of various hygienic and biosecurity practices essential for minimizing the risk of Salmonella transmission. Similar findings were reported where meat sellers exhibited a little higher than moderate concern for meat hygiene [27]. In another study, positive attitudes of the participants indicating the willingness among to engage in proper practices while knowledge may be lacking [32]. A particularly strong consensus was observed around avoiding face and hair contact during meat processing (ES=0.16). This may be attributed to visible and culturally reinforced perceptions of cleanliness during food handling, which are often reinforced by peer practices and consumer expectations in traditional marketplaces [21]. Similarly, the recognition of training as essential to reducing Salmonellosis risk (ES=0.13) is noteworthy. This finding aligns with previous studies emphasizing that carcass handlers’ express willingness to training for improving the food hygiene and sanitation [42]. The sellers’ positive attitude toward hand hygiene, such as avoiding money handling during meat processing (ES=0.10), reflects an encouraging understanding of indirect contamination routes. These might be due to a considerable proportion of vendors (47.6 %) believed that risk-free dressing and processing of chicken without having contamination is a part of the vendor’s duties [14]. Moderate but significant agreement was observed for using clean water and maintaining sanitary environments during meat dressing. In other study, 77% of the meat retail seller were agreed to keeping sanitary environment is difficult [27]. Particularly, the awareness of environmental hygiene, likely influenced by consumer demand and increasing media coverage on foodborne diseases [20]. The emphasis on offal disposal highlights the recognition of waste management as part of infection control a topic rarely addressed in informal market training but increasingly spotlighted in community-based awareness campaigns. Interestingly, sellers showed more divided opinions regarding personal grooming practices, such as avoiding long nails. While 32.64% moderately agreed and 26.16% strongly agreed with this hygiene aspect, the comparatively lower agreement suggests a potential gap in the understanding of how personal cleanliness contributes to cross-contamination. In another study about 70% of the meat seller believed that their nail should be small [27]. The male sellers, those with higher education levels, and participants who had TLPM exhibited significantly more positive attitudes. This reinforces earlier findings that education enhances critical thinking and risk perception regarding zoonoses [43]. The strong predictive value of higher education in both univariate and multivariate logistic regression models (Bachelor’s degree holders had an OR of 13.48) underscores the transformative impact of formal education on fostering responsible food handling attitudes. In order to improve knowledge and adherence to sanitary requirements and guarantee the biosecurity of raw poultry meat, it might be helpful to understand the subtle differences in attitudes across various occupational groups [27,28]. Further study into the elements that influence attitudes regarding meat shop biosecurity practices by these groups might give a more thorough knowledge of the dynamics at play.

This study sheds light on the biosecurity practices adopted by poultry/poultry-meat sellers in relation to the risk of zoonotic salmonellosis, revealing a mix of encouraging behaviours and critical gaps. In the current study, the practice of separating poultry by age, type and sickness was reported by 36.34% of sellers as “always” followed, indicating a basic understanding of disease prevention principles. Segregation of poultry is a recognized strategy in reducing pathogen transmission across groups [44]. This findings are also similar to other study [45] where ≥95% of commercial poultry farm in Rajshahi of Bangladesh separated (isolation) the sick birds. Particularly, poultry/poultry-meat sellers keep the live poultry for short period of time due to sell. However, the relatively low proportion of consistent application, along with small effect sizes, suggests that while awareness may exist, consistent execution is lacking. Mostly it was observed in earlier studies that the general awareness of meat handlers or vendors were always positive but they had relatively poor practices [14,22]. Hand hygiene, a key intervention in preventing zoonotic disease transmission [46], was reported by 38.19% of sellers as being practiced “every time” before and after meat processing. This figure is moderately encouraging compared to similar settings, where studies have found that 42.8% chicken meat sellers wash their hands before meat processing and 87.2% wash after meat processing [14]. These differences might be due to fluctuation of practices. In Ethiopia, a study reported that 68.8% of respondents didn’t wash their hands after coming into contact with animals [2]. However, the inconsistent use of PPE, such as masks, aprons, and gloves with only about 15% always using them remains a serious concern. Similar findings also reported in another study in Bangladesh where only 30 % of respondents used face masks and 14 % used caps during meat processing and selling [27]. Proper PPE use is a cornerstone of occupational safety in meat handling environments, and its limited adoption among sellers might increases the chance of potential zoonotic pathogens transmission like as *Salmonella spp*. Another critical area is waste management and environmental hygiene. A considerable portion of sellers reported the consistent use of designated dustbins for slaughter waste and hygienically disposed the dead birds and slaughtering waste. The open dumping of poultry wastes significantly affects the surroundings including the heavy metal leaching, greenhouse gas emissions, potential breeding sites of mosquitoes that in turn could lead to serious public health issues [47]. In the poultry shops, regular use of fresh (14.81%) and hot (36.57%) water for dressing of poultry meat, washing of utensils (e.g., knife, chopping board etc.; 18.75%) and cleaning the shop with disinfectants (33.56%) that also reflects moderate adherence to best hygienic practices. These hygienic practices are very important, because the cross-contamination from raw meat to different foodstuffs and utensils is a primary source of foodborne illness outbreaks [48]. Moreover, dressing by hot water along with fresh water is effective, because the viable Salmonella counts of are relatively not affected when using only normal water [38]. In a study in the capital city of Bangladesh reported that 50.3% wash knife and 51.7% wash their chopping board before use where 46.6% of chicken vendors clean their shop with disinfectant at the time of closing [14]. These findings are somehow difference from our study and it might be due to variation in the study area. However, these practices are essential in limiting the spread of pathogens. Moreover, regular use of disinfectant is crucial for poultry biosecurity practices in Bangladesh [12,49]. In this current study, a considerable proportion of poultry/poultry-meat sellers showed unexpected personal habits practice (eating, drinking, and smoking inside the shop and wearing jewelleries during work) which is consistent with other studies [14,18,21]. However, control of external contamination sources such as dogs, cats, and wild birds was less frequently practiced, despite being critical in controlling zoonotic risk from environmental reservoirs. Moreover, control of these external contamination sources is effective to minimize the spread of infectious diseases of poultry [12,49]. The demographic associations further enrich the understanding of factors influencing these practices. Male sellers demonstrating significantly higher correct biosecurity practice scores than females. This might be due to male dominance in physically demanding market roles and increased access to training or peer networks with better compliance. Our findings shows that young of 26 to34 year age were involved in good practices though the significant proportion of older (≥55 years ages) had good knowledge. These findings indicate, tough the older may have good knowledge but more numbers of youngers are involved in correct biosecurity practices. In Lebanon, the participants aged between 43 and 55 years were more likely (OR =9.931, p<0.05) involved in good practice levels compared to those aged 18–29 years [32]. In another study in Bangladesh reported that middle-aged (31 to 45 years) meat sellers have the highest mean of practices score (13.73) in meat hygiene practices [14]. Our findings suggest that educational level had emerged as a strong predictor of biosecurity practices for zoonoses control. Accordingly, sellers with higher secondary or bachelor’s education were significantly more likely to adopt correct practices, which is consistent with other studies [14]. Similar reports were also observed in Ethiopia, where participants with secondary education or higher had more tendency (OR= 6.76, p<0.05) to correct practice against zoonoses transmission [2]. Education may enhance cognitive abilities to interpret zoonotic risks and promote rational decision-making in daily practices. Training-related variables, such as TLPM and TPMPM, were significantly associated with better biosecurity practices in the univariate analysis (OR=2.33 and OR=1.67 respectively), highlighting the importance of training programs [21] to enhance the biosecurity practices among the poultry/poultry-meat sellers in Bangladesh. However, these associations lost significance in the multivariate model, suggesting that education may mediate the effect of training. Similar to our findings, the slaughterhouse workers and retail meat sellers who received training showed that the good practices score (13.46) compared to those who (12.05) did not in Bogura District of Bangladesh [27]. The study in Morocco also reported that the of training on regarding meat hygiene, safety during carcass processing is essential for good slaughtering practices [42]. Moreover, other study reported that individual without the biosecurity training were significantly (p<0.01) more likely (OR=4.80) to engage in zoonotic disease-related practices in Bangladesh [25]. Perhaps training are the major sources of information about zoonotic diseases.

The present study revealed statistically significant positive correlations between knowledge, attitudes, and practices (KAP) scores among poultry and poultry meat sellers, underscoring the interconnectedness of these domains in shaping biosecurity behaviours. Different studies on KAP also reported the significant positive correlations among KAP score [19,21]. The positive correlation between knowledge and attitudes (r =0.221, p<0.01) suggests that individuals with greater understanding of zoonotic salmonellosis can shape the perceptions of risk and the importance of preventive actions. However, the relatively low coefficient also implies that knowledge alone may not be sufficient to drive strong attitudinal shifts. Factors such as cultural beliefs, misinformation, or perceived constraints may moderate this relationship. A low strength correlation was found between knowledge and practices (r=0.281, p<0.01), reinforcing the idea that higher knowledge scores lead to improved biosecurity practices related to handling poultry and raw poultry meat [14,21,27,42,50]. Nevertheless, the low strength of the correlation indicates that knowledge must be translated into actionable behaviour through supportive structures, such as training, incentives, or regulatory enforcement. The moderate strength relationship observed between attitudes and practices (r=0.371, p<0.01), suggesting that sellers with positive attitudes toward biosecurity measures and zoonotic risks prevention are more likely to engage in protective practices and behaviours of sellers [24]. Perhaps, this emphasizes the critical role of fostering positive attitudes to achieve better compliance with recommended practices.

Sellers with good knowledge scores rated their perception of zoonotic risk significantly higher, with a median Likert score of 7. This indicates that knowledge plays a pivotal role in shaping perceived susceptibility and seriousness of disease threats. This finding supports that awareness of disease transmission enhances perceived vulnerability and motivates individuals to engage in preventive actions [51]. Similarly, sellers with positive attitudes and correct biosecurity practices also showed significantly higher perceived risk scores. This alignment of perception indicating the individuals who recognize zoonotic threats are more inclined to adopt protective behaviours [26]. The correlation analysis further underscores the importance of demographic variables in shaping perceptions. Education and TLPM were significantly associated with higher PPRZDT, indicating that sellers with greater access to formal education and training are more aware of zoonotic disease risks [2,14,21,25,27]. Gender was found to have a significant correlation with PCPBM. This finding may reflect cultural and role-based differences in how male and female sellers view their ability to control risks. Importantly, the lack of significant correlations of demographic variables with PIZETP suggests that interest in learning more about zoonoses may be uniformly distributed across seller profiles.

## Conclusion

This research reveals that the few live poultry/poultry-meat sellers realized Salmonella in meat is a zoonotic disease and recognized cleanliness matters, many still did not clearly understand how to avoid contamination and how to follow hygiene practices consistently. Someone’s opinion about managing risk was usually optimistic, especially if they had more schooling and past experience. Still, the application of required practices such as using personal protective gear and keeping the environment hygienic, was not reliably carried out. Study participants’ education level, age and earlier training had the strongest link to their knowledge and attitudes and level of education was a greater factor affecting biosecurity behavior than income or training history. Significantly, our findings showed that attitudes related more strongly to actual actions than knowledge did, suggesting we should pay equal attention to mindset and information. Despite this, sellers who were better educated and training often believed zoonotic disease was risky, but most sellers, regardless of education, rated themselves as confident in their biosecurity and wanted to learn more in training programs. As a result, everyone seems willing and eager to learn, so education programs can make the most of this by providing practical and inclusive lessons. The findings suggest that training programs should be adapted for different places, helping people develop confidence, practice needed skills and stick to good hygiene and safety habits in the long run for biosecurity measure to control the zoonoses.

## CRediT authorship contribution statement

**Mirza Mienur Meher:** Conceptualization, Resources, Writing-original draft, Writing-reviewing, Visualization, Validation, Methodology, Project administration, Formal analysis, Investigation, Data curation. **Marya Afrin:** Resources, Writing-reviewing, Validation, Methodology, Project administration. **Abdullah Al Bayazid:** Investigation, Methodology, Data curation. **Nashib Parajuli:** Investigation, Methodology, Data curation. **Md Sohel Rahman:** Investigation, Methodology, Data curation. **Md Sayedul Islam:** Validation, Methodology **Md Zulfekar Ali:** Validation, Methodology. **Md. Mominul Islam:** Validation, Methodology.

## Acknowledgement

The authors would like to express gratitude to the participating poultry/poultry-meat sellers for voluntarily taking part in the survey despite of their busy schedule. The authors also would like to thank all the persons who assisted during the data collection process.

